# Loops, TADs, Compartments, and Territories are Elastic and Robust to Dramatic Nuclear Volume Swelling

**DOI:** 10.1101/2021.08.20.457153

**Authors:** Jacob T. Sanders, Rosela Golloshi, Peyton H. Terry, Darrian G. Nash, Yang Xu, Job Dekker, Rachel Patton McCord

**Author notes:** These authors contributed equally to this work (Jacob T. Sanders, Rosela Golloshi). Corresponding author: Rachel Patton McCord; 309 Ken & Blaire Mossman Bldg. 1311 Cumberland Ave. Knoxville, TN 37919.

## Abstract

Layers of genome organization are becoming increasingly better characterized, but less is known about how these structures respond to perturbation or shape changes. Low-salt swelling of isolated chromatin fibers or nuclei has been used for decades to investigate the structural properties of chromatin. But, visible changes in chromatin appearance have not been linked to known building blocks of genome structure or features along the genome sequence. We combine low-salt swelling of isolated nuclei with genome-wide chromosome conformation capture (Hi-C) and imaging approaches to probe the effects of chromatin extension genome-wide. Photoconverted patterns on nuclei during expansion and contraction indicate that global genome structure is preserved after dramatic nuclear volume swelling, suggesting a highly elastic chromosome topology. Hi-C experiments before, during, and after nuclear swelling show changes in average contact probabilities at short length scales, reflecting the extension of the local chromatin fiber. But, surprisingly, during this large increase in nuclear volume, there is a striking maintenance of loops, TADs, active and inactive compartments, and chromosome territories. Subtle differences after expansion are observed, suggesting that the local chromatin state, protein interactions, and location in the nucleus can affect how strongly a given structure is maintained under stress. From these observations, we propose that genome topology is robust to extension of the chromatin fiber and isotropic shape change, and that this elasticity may be beneficial in physiological circumstances of changes in nuclear size and volume.

## Introduction

Imaging and chromosome conformation capture-based approaches have revealed that chromosomes are organized at different length scales into loops, topologically associating domains (TADs), compartments, and chromosome territories (Gibcus and Dekker 2013; Dekker and Mirny 2016; McCord et al. 2020). Mechanisms such as loop extrusion, phase separation, and tethering of chromatin to nuclear structures are emerging as key players in building and maintaining these chromosome structures (Fudenberg et al. 2016; Strom et al. 2017; van Steensel and Belmont 2017; Ganji et al. 2018; Nuebler et al. 2018; Golfier et al. 2020). However, less is known about the physical properties of these structures and how they respond to perturbation. Are certain features of genome structure held together specifically and robustly while others are weaker or an incidental effect of nuclear crowding? Carefully quantified disruption of isolated nuclei has proven to be a fruitful approach to measure properties of chromosome structures. Fragmenting chromatin inside intact nuclei to varying degrees has revealed the relative stability of different types of chromosome interactions, from strong lamina associations to more dynamic polycomb associated interactions (Belaghzal et al. 2021). But, other approaches are needed to measure the elasticity and stability of these interactions in the context of the intact chromosome polymer.

A variety of approaches have been used over the past decades to investigate the physical properties of nuclei and chromosomes, and the results suggest that the 3D genome structure has some elastic properties and some potential for irreversible deformation. When isolated nuclei are stretched by physiological-level forces, the chromatin structure acts like a stretched spring that provides resistance to initial deformations as it stretches, with a spring constant that varies with altered histone modifications and chromatin compaction (Stephens et al. 2017a). Conversely, chromatin also has some properties of a compressed spring. Disrupting interactions between histone tails dramatically releases the energy that is stored in this ‘prestressed’ compressed state of genome folding, even to the point of exploding the nucleus (Mazumder et al. 2008). Micropipette aspiration experiments have shown that chromatin domains stretch, sometimes irreversibly, as they are pulled into a narrow opening (Irianto et al. 2017). Microscopic observations have demonstrated visible changes in chromatin structure inside the nucleus after external forces are applied to cells (Guilak 1995; Maniotis et al. 1997), and such chromatin deformations can even directly induce gene expression in live cells (Tajik et al. 2016). In all these experiments, however, chromosomes are typically conceived as coarse-grained objects with uniform physical properties, and there has been little investigation into how specific genomic regions or structures either change or maintain their organization during deformation.

External force exertion on the nucleus likely affects regions of the genome near the applied force disproportionally to the rest of the genome. Thus, localized external nucleus stretching will not allow us to characterize the response of the genome structure to physical perturbation genome-wide in a population of cells. So, in this work, we use low-salt chromatin expansion: rather than pulling on the exterior of the nucleus, salt depletion changes the charge balance along the chromatin, expanding the local chromatin fiber itself, and thus expanding the nucleus from the inside out. This will allow us to monitor the effects on 3D genome topology chromatin fiber extension and nucleus expansion.

Low-salt swelling of isolated chromatin fibers or nuclei has been used for decades to investigate the structural properties of chromatin. Treatment of intact nuclei with EDTA to chelate magnesium ions enabled electron microscopic images of networks of chromatin fibers, but with little ability to interpret the structural patterns observed (Woods et al. 1991). Electron microscopic studies of isolated chromatin fibers concluded that removing magnesium and sodium ions from the chromatin causes a state change from a 30 nm fiber to an extended 10 nm fiber (Widom 1986). Similarly, isolated mitotic chromosomes undergo reversible decondensation and a decrease in stiffness in low-salt conditions, again, presumably explained by a change from 30 to 10 nm fiber (Poirier et al. 2002). However, the simple idea of a conversion between a 10 and 30 nm fiber does not explain what would happen to the layers of 3D genome structure in an interphase nucleus during such swelling. In fact, recent work suggests that the 30 nm fiber is uncommon in the *in vivo* interphase genome structure (Dekker 2008; Razin and Gavrilov 2014; Maeshima et al. 2016; Ou et al. 2017; Maeshima et al. 2019; Krietenstein and Rando 2020), so further work is needed to understand how such *in vitro* studies translate to the interphase nucleus. Imaging of swollen nuclei after photobleaching has previously suggested that the large scale chromatin structures return to their initial positions after swelling and return to the original nucleus size (Mazumder et al. 2010b). Meanwhile, experiments with DNA-damaging agents have suggested that notable decondensation of the chromatin occurs during low-salt swelling: chromatin in swollen nuclei is more susceptible to DNA damage by irradiation than high-salt compacted chromatin (Takata et al. 2013). So, the question remains, what structures are decondensing during swelling? Is the whole 3D structure really returning to the same state after re-condensation as suggested from whole nucleus observations or are there more subtle effects at a small scale?

In this work, we combine the classic low-salt swelling perturbation approach with detailed measurements of 3D genome interactions using Hi-C. By doing so, we can probe how the layers of genome structure react to or maintain robustness to this physical perturbation. At the same time, similar to expansion microscopy approaches (Chen et al. 2015), we may be able to detect connections in the structure more clearly by spreading out genomic regions that are not linked by stable interactions.

## Results

### Chromatin expansion is visibly elastic

To avoid the complicating effects of secondary biological signaling cascades on 3D genome organization, we performed all our experiments with isolated human nuclei. As reported previously (Takata et al. 2013), we can expand isolated human lymphoblast GM12878 nuclei by replacing a buffer containing physiological levels of cations sodium and magnesium (1x HBSS) with a buffer with no cations and EDTA to chelate any remaining magnesium (Figure 1A). Expansion happens instantaneously (Movie S1), indicating that this expansion is a purely physical perturbation, not requiring biologically controlled signals. This initial rapid expansion is followed by a slower further increase in nucleus size over the next 10 minutes to 1 hour (Figure 1B). When a nucleus is returned to 1x HBSS, it shrinks back to its original size, again effectively instantaneously (Movie S2). There is variability in the precise extent of expansion of each nucleus within the population, but all nuclei show at least some expansion, with the average expansion being 2-5 fold in cross-sectional area or 4-10 fold in volume (Figure 1B,C). We find that staining chromatin with DAPI causes a measurable shrinking of the expanded nucleus (Figure S1A,B). So, we rely on phase contrast light microscopy to measure the extent of nucleus expansion. However, staining chromatin with DAPI demonstrates that the chromatin fills the entire space of the expanded nucleus (Figure S1A), and it is not the nuclear membrane alone that expands.

**Figure 1.**
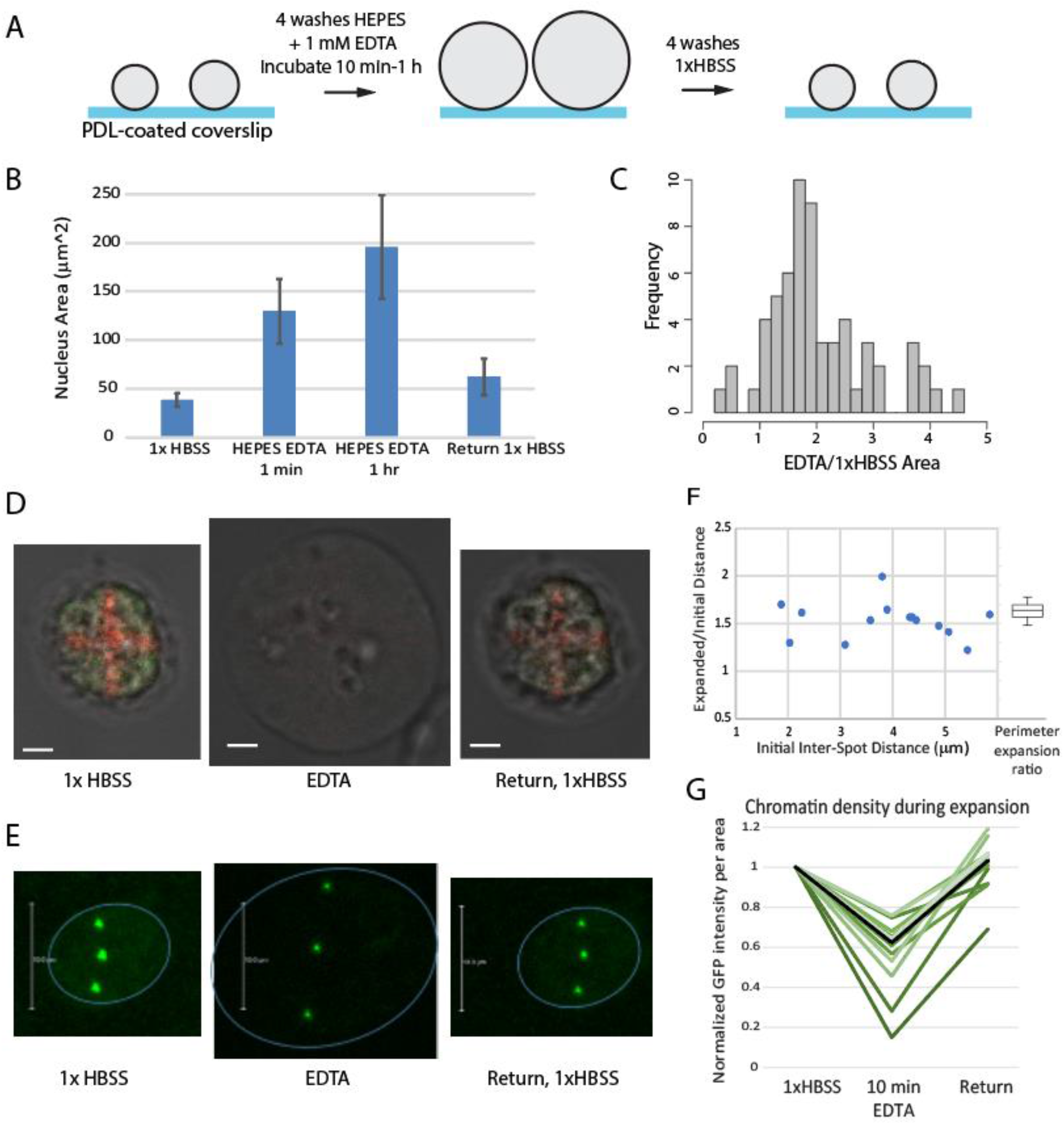
Elasticity of nuclear organization is evident in single cell imaging. A) Experimental design. B) Mean cross-sectional area changes of GM12878 nuclei before, during and after expansion in EDTA (N = 30; error bars = std. dev.) C) Distribution of fold changes in area of individual GM12878 nuclei immediately after EDTA addition (N = 61) D) Photoconverted pattern on GM12878 nuclei transfected with pDendra2-H4 is still visible when nuclei are returned to 1xHBSS after expansion in EDTA (scalebar = 2 mm), nucleus treated with RNase prior to experiment. E) CRISPR-dCas9-GFP labeling of pericentromeric repeats of chr3 in U-2 OS cells. Chromosomes spread out during expansion in EDTA but return to their original locations after return to 1x HBSS. Blue lines indicate location of nuclear periphery. Scalebar = 10 mm F) The ratio of expanded vs. initial distances between each pair of chr9 spots in HEK293T cells (y axis) is independent of initial distance between spots and is comparable to the fold increase in perimeter of these same cells (boxplot, right) G) Quantification of GFP intensity per area during expansion of CRISPR-GFP labeled chr9 repeats in HEK293T cells. Green lines = intensity trajectories of individual chr9 spots; Black line = median of all intensity measurements.

We first evaluated the previously reported elasticity of the genome structure by monitoring a photoconverted pattern on nuclei during expansion and re-contraction. Histone H4 fused to photoconvertible Dendra2 was expressed throughout the nucleus and then photoconverted in a specific pattern by 405 nm laser light, prior to expansion. Upon expansion, the fluorophores are spread out such that the pattern is barely visible, but, when the nucleus is recontracted by the replacement of salt, the same pattern that was present before expansion was visible again (Figure 1D). The photoconverted pattern was also restored upon recontraction in other cell types, and when nuclei were left in the expanded state for an hour (Figure S1C,D) or treated with RNase (Figure S1E).

The visibly apparent elasticity of nucleus expansion is also observed in nuclei with mutant Lamin A and after RNA degradation. Cells expressing progerin, the mutant form of Lamin A that causes Hutchinson-Gilford Progeria Syndrome (HGPS), have stiffer nuclei with characteristic blebs (Shumaker et al. 2006; Ribeiro et al. 2014), and previous work has suggested that these altered mechanical characteristics make progerin expressing nuclei unable to recontract after expansion (Dahl et al. 2006). In contrast, we find that fibroblast nuclei stably expressing progerin are able to expand and re-contract (Figure S2A and B). Visible wrinkles in the lamina spread out and become smoother during expansion, but then return to nearly identical positions after contraction. This result held true whether we used transient transfection of progerin in a different cell line (HeLa), progeria patient cells, or used alternative low and high salt buffer conditions for expansion.

### Labeling specific loci allows more detailed quantitation of chromosome expansion

These results show a generic elasticity of the genome structure, but do not provide information about the positions of any specific genomic loci. To investigate the effects of expansion on the position of specific chromosomes, we took advantage of the CRISPR imaging system for targeting fluorophores to genomic loci using catalytically inactive Cas9 and sgRNAs complementary to chromosomal repeat regions (Ma et al. 2015). Transfection efficiency of these large plasmids was very low in GM12878 cells, so for these experiments, we used U2-OS and HEK293T cells that are easier to transfect. Due to the abnormal karyotype in these cells, 3 fluorescent spots are observed, representing the pericentromeric regions of 3 copies of chromosome 3 or chromosome 9 in these cells. These pericentromeric chromosome repeat regions spread out from each other proportionally to the swelling of the entire nucleus, and then return to their original positions upon contraction (Figure 1E, F, Figure S3A).

We further noticed a loss of fluorescence intensity at individual loci in the expanded state, which increased again after re-contraction. After quantifying the fluorescence intensity during expansion for 13 individual loci, we found that all foci decreased in intensity during expansion and recovered during contraction, though with a notable amount of variability in how much each spot increased and decreased. The intensity loss per area is consistent with the cross-sectional area of the chromosome region increasing 1.3-6.6 fold, consistent with the range of increase in size measured for whole nuclei.

The median intensity loss during expansion was 40%, and on average the fluorescence recovered to 100% after re-contraction (Figure 1G). This change in fluorescence intensity is corrected for the intensity losses caused by repeated imaging, and cannot be explained by a dissociation and re-association of fluorescent proteins, as demonstrated by the lack of any fluorescence recovery after local photobleaching of a labeled locus (Figure S3B). Thus, we conclude that the decreased fluorescence per volume during expansion reflects a local decondensation of the labeled peri-centromeric repeat chromatin region in low-salt conditions. Overall, we can observe that reversible local and global spreading of loci occurs with this expansion.

### Hi-C experiments reveal remarkable preservation of genome contacts in expanded nuclei

While microscopy shows quantifiable decondensation and spreading out of labeled genomic loci during expansion, it cannot reveal how specific chromosome structures are affected by this perturbation. To investigate the effect of low-salt nuclear swelling on 3D genome topology and contacts, we performed Hi-C experiments in the 1x HBSS, EDTA, and return to 1x HBSS conditions according to established protocols (Belton et al. 2012), with modifications necessary to adapt the protocol for adherent isolated nuclei (see Methods). GM12878 nuclei were crosslinked directly on the dishes in which expansion took place, and image quantitation indicates that the expanded size of the nuclei was preserved during this fixation (Supplementary Figure S3C). The Hi-C protocol utilized was more similar to “dilution” rather than “in situ” Hi-C because nuclei lysis before digestion and ligation was necessary to recover crosslinked chromatin from the poly-D-lysine dishes. As expected, this led to more variable cis/trans ratios among replicate experiments (Table S1) (Nagano et al. 2015; Golloshi et al. 2018). In each replicate set, the expanded nuclei EDTA condition was more divergent while 1xHBSS and the return to 1xHBSS conditions were most highly correlated (Figure S4A). We present detailed analyses of the highest quality and most deeply sequenced replicate, while checking overall conclusions in the other replicates (Figure S4).

After mapping and correcting the Hi-C data for inherent biases using iterative correction (Imakaev et al. 2012), we examined the contact maps at various resolutions, from the whole genome scale down to the level of individual loops. Our results indicate a remarkable preservation of genome structure at all length scales.

At the scale of chromosome territories, Hi-C results indicate that both the enrichment of interactions within individual chromosomes and the patterns of neighboring chromosome arrangements are preserved during nucleus expansion (Figure 2A and B). The population average evidence for the preservation of chromosome territories during expansion was also validated at the individual nucleus level by multicolor fluorescence in situ hybridization (mFISH) (Figure 2C), though the volume of the expanded nuclei was artificially reduced by the hybridization procedure necessary for FISH, as noted previously (Woods et al. 1991). These results indicate that expansion does not cause randomization of chromosome territory locations or significantly more intermingling of chromosomes. The pattern of compartmentalization of genomic regions into the active (“A”) and inactive (“B”) compartments is also remarkably preserved during expansion, as shown by the highly similar plaid interaction pattern and first principal component of an individual chromosome contact map (Figure 2D,E, Figure S4B).

**Figure 2.**
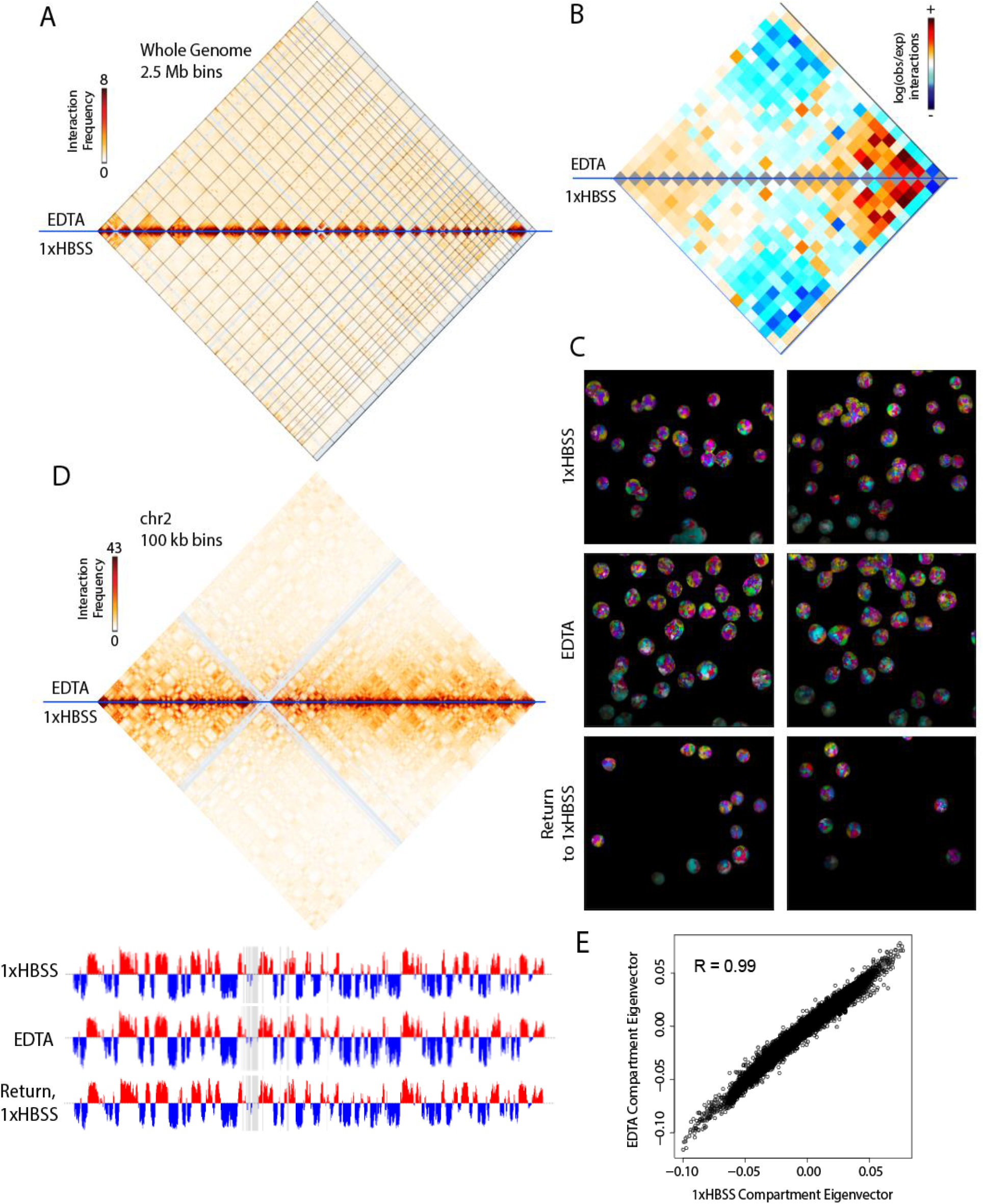
Major features of 3D genome topology are preserved during nuclear expansion. A) Genome-wide contacts from Hi-C performed on GM12878 in EDTA expansion (top) and 1xHBSS (bottom) in 2.5 Mb bins. Black lines = divisions between chromosomes, arranged left to right (chr1-22, X). B) Observed vs. expected contacts between each pair of chromosomes, left to right chr1-22, X. C) Multicolor FISH probes indicate the persistence of chromosome territories through expansion and contraction. D) Contacts within chr2 in 100 kb bins and the first principal component (PC1) below shows little change between A (red) and B (blue) compartmentalization. E) PC1 values are highly correlated between expanded and non-expanded conditions in all 100 kb bins genome wide.

At somewhat higher resolution (40 kb bin size), we observe a similar segregation of genomic regions into topologically associating domains (TADs) in both original and expanded conditions (Figure 3A). The insulation score profile is calculated as previously described using interaction sums within a sliding window along genomic coordinates (see Methods, (Crane et al. 2015)). Dramatic dips in this profile correspond to TAD boundaries, and the depth of these dips indicates the degree of contact insulation and thus represents the strength of the TAD boundaries. Overall, this insulation profile is highly similar between all three conditions, as can be observed in an example 10 Mb region (Figure 3A) and quantitatively across whole chromosomes (Figure 3B, Figure S4C). At 40 kb bin resolution with a 520 kb bin insulation square size, there is little change in the average insulation profile across all TAD boundaries on either large, gene poor (chr2) or small, gene dense (chr19) chromosomes (Figure 3C).

**Figure 3.**
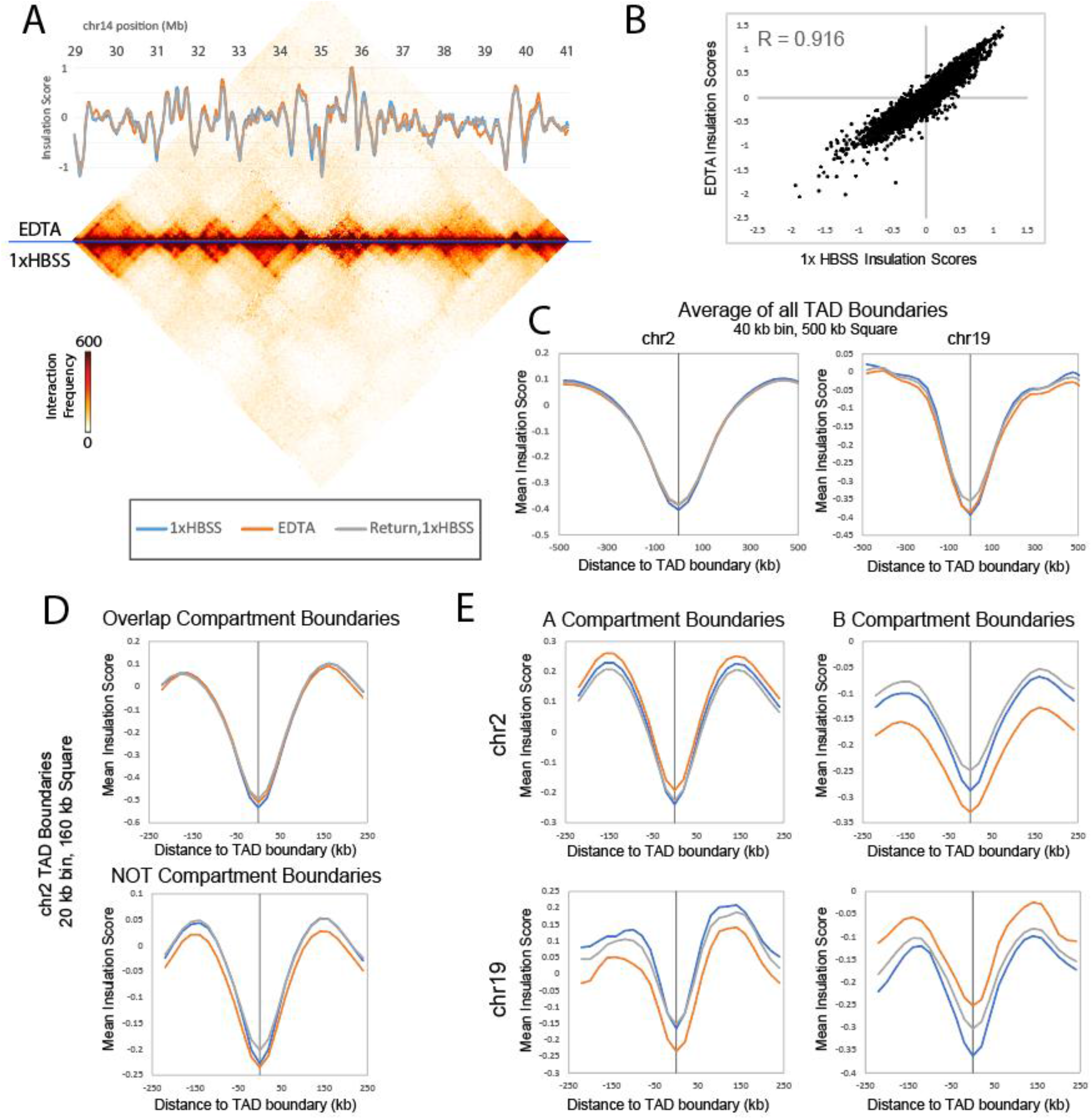
TADs are generally preserved during nuclear expansion. A) Contact map across 12 Mb of chr14 shows the general similarities and minor differences between 1xHBSS (bottom) and EDTA (top). Insulation score (20 kb bin resolution, 160 kb insulation square) is shown overlaid with the heatmap. B) Insulation scores (calculated same as A) are well correlated (chr2 shown) between expanded and original size nuclei. C) At 40 kb bin resolution with 520 kb insulation square size, little change seen in average insulation profile at TAD boundaries. D) Average insulation profile at higher resolution (20 kb bin, 160kb insulation square) around TAD boundaries that either do (top) or do not (bottom) overlap compartment boundaries. E) Average insulation for TAD boundaries in the A or B compartment on chr2 or chr19.

However, when boundaries are evaluated at higher resolution and in specific categories, region-specific changes with expansion start to emerge. We observe that TAD boundaries which overlap with compartment boundaries do not show alterations in insulation profile (Figure 3D), echoing the previous observation that compartmentalization is not changing with expansion. However, TAD boundaries that occur within compartments do show an apparent change in strength during expansion (Figure 3D). We find that this overall change is actually the sum of two opposing changes in different types of chromatin. On chromosome 2 (and other large, gene poor chromosomes) TAD boundaries in the A compartment experience an overall upward shift of insulation score while boundaries in the B compartment experience a downward shift in insulation score during expansion (Figure 3E). Upon return to 1x HBSS, the insulation profile approaches a return to the original state, though with slightly weakened boundaries overall. Strikingly, on chromosome 19, we find the opposite effect, where A compartment boundaries experience a decrease in insulation score while B compartment boundaries experience an increase in this profile (Figure 3E).

This opposing behavior of chr2 and chr19 likely relates to a difference in chromatin types between chr2 and chr19. In terms of subcompartment types (Rao et al. 2014), chr19 has only the A1 active compartment type while chr2 is predominantly (80%) A2 type in its active regions. Meanwhile, chr2 heterochromatin is predominantly (70%) B3 type, which is characterized by lamin associated regions, while chr19 has no B3 type chromatin and instead has a majority of nucleolar-associated regions (NADs). Chromosome 19 has few lamin associated domains (LADs) while chromosome 2 has many (Kind et al. 2015). Correspondingly, chr19 tends to be located in the interior of the nucleus of GM12878 cells while chr2 is located at the periphery (Tanabe et al. 2002; Das et al. 2020). Indeed, when the changes in insulation profile are considered around boundaries in specific sub-compartments, we observe that chr19 B1 (polycomb-related) regions behave more like chr2 B compartment regions while chr19 B4 NAD regions match the insulation change seen in chr19 B compartment regions overall (Figure S5). Importantly, these overall shifts in the insulation profile, rather than relating only to the TAD boundary segregation, suggest an overall change in the local scaling of contacts with distance during expansion, so next, we turned to examination of this contact scaling.

### Contact scaling changes reflect elongation of the local chromatin fiber during low-salt expansion

Plotting the average interaction frequency over distance separating two genomic loci can reveal properties of the chromosome fiber (Fudenberg and Mirny 2012). At 40 kb resolution, considering all intrachromosomal interactions across all chromosomes, the scaling plot of interactions over distance shows higher interaction frequency at large genomic distances for both expanded and return to 1x HBSS conditions (Figure 4A). This suggests that at a large scale, expansion of the chromosomes results in more intermingling of distant chromosome regions, and that some degree of intermingling persists upon return to a normal nucleus size. At smaller distances, a steeper drop in interactions observed in the expanded condition only (Figure 4B, Figure S4D). This measurement captures the extension of the local chromosome fiber previously seen in electron microscopy (Widom 1986) as the low cation condition alters the charge shielding between nucleosomes. Motivated by the different insulation results observed for different types of chromatin above, we constructed separate scaling plots for A and B compartment regions on chr18 (gene poor, peripheral, LAD-rich) and chr19 (gene dense, interior, NAD-rich). We observe that the local interaction decrease with expansion occurs in both A and B compartments, but A compartment interactions change more on chr19 than on chr18 (Figure 4C).

**Figure 4.**
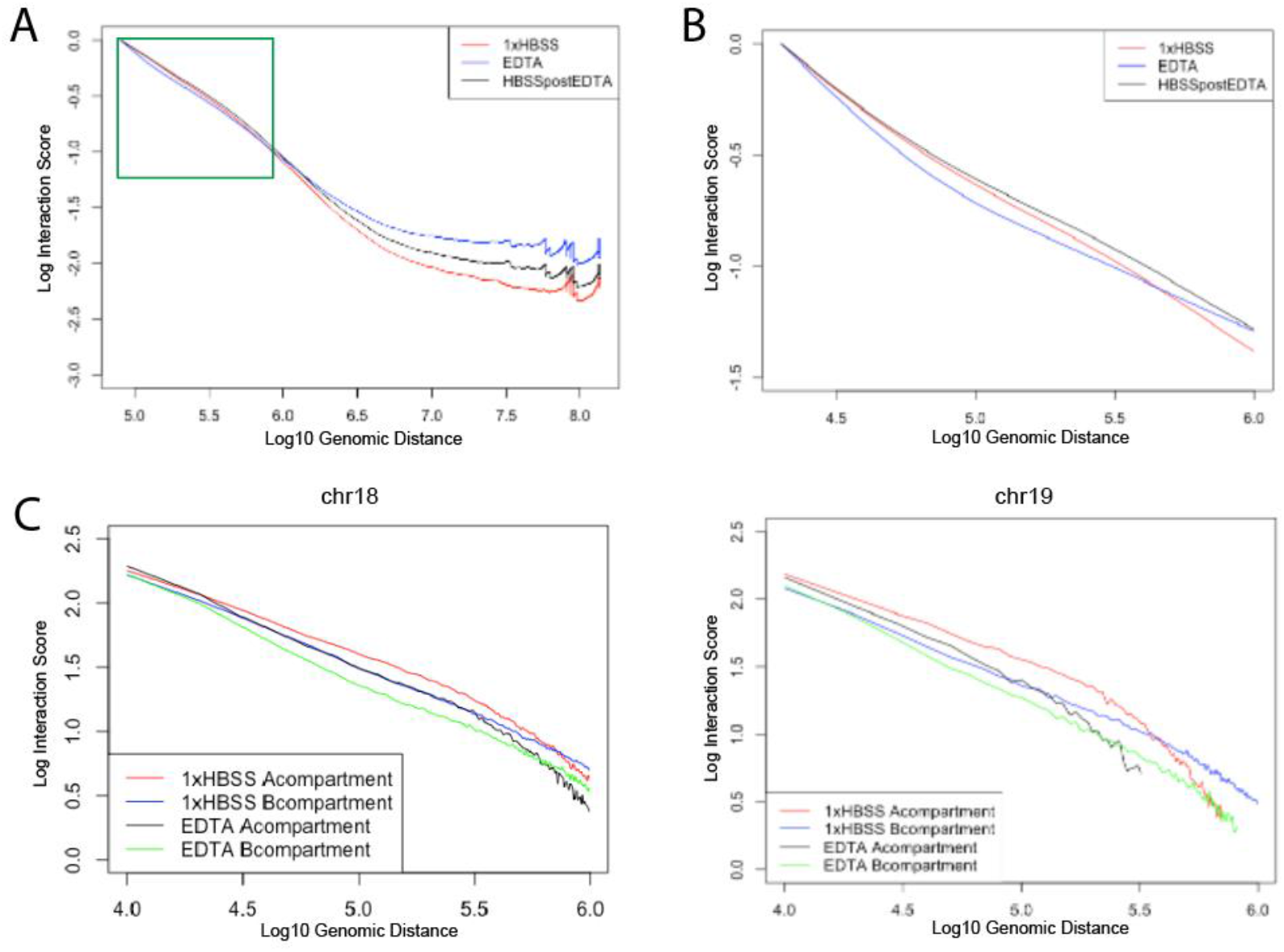
Polymer scaling is altered reflecting chromatin fiber elongation during expansion. A) Log10 interactions vs. Log10 of genomic distance in base pairs for all 40 kb binned intrachromosomal interactions genome wide. A faster drop in local interactions is seen at short distances in EDTA only (green box). B) 10 kb resolution interaction scaling genome wide, zooming in to interactions within 1 Mb. C) Interactions vs. distance separately for regions in the A or B compartment on chr18 or chr19.

### Differential changes in 3D genome structure with nucleus position and lamin association

While the A or B compartment assignment of genomic bins across the genome changes very little during expansion, we do observe a quantitative loss of interactions within compartments in the expanded state (Figure 5A). Interestingly, the loss of interactions is strongest within the B compartment on small, gene rich interior chromosomes, (chr19) while interactions within the B compartment are more preserved in chromosomes located near the nuclear lamina, such as chr2 and chr18. We already saw above that the overall decay of interactions with distance changed differently in different chromosome regions. We can quantify this more precisely by plotting a ratio of distal to local interactions (DLR) in each bin across the chromosome (Heinz et al. 2018). This DLR measure quantifies how much each 100 kb genomic bin interacts with distal (greater than 3 Mb away) or local (closer than 3 Mb in linear genomic distance) regions along the chromosome and is related to the LOS metric used to quantify changes in Hi-C interactions after chromosome fragmentation (Figure 5B) (Belaghzal et al. 2021). On gene poor chromosomes at the nuclear periphery (e.g. chr2 and chr18), B compartment regions show an increase in distal relative to local interactions in the EDTA expanded nucleus (Figure 5C). Gene dense chromosomes such as chr19 do not follow the same trend, however. This again echoes the finding first observed for TAD insulation analysis that different types of chromosome regions behave differently during expansion. These different responses may be explained by differences in lamin or nucleolar associations of each chromosome. Genome wide, regions classified as constitutively associated with the nuclear lamina (cLADs) (Kind et al. 2015; de Graaf et al. 2019) show the strongest increase in distal vs. local interactions during expansion (Figure 5D, Figure S4E). The chr19 B compartment is composed of facultative LADs (fLADs), rather than cLADs, and this type of region does not have a strong increase in DLR with expansion. Both chr17 and chr19, which have few LAD domains and instead strong nucleolar associations, show a different pattern of interchromosomal interaction change upon expansion (Figure 5E). While chr18 shows an increase in its interactions with other chromosomes upon expansion, chr17 and 19 interact more within themselves and less with other chromosomes after expansion. These results imply that lamin associations hold more distal regions of chromosomes together during the general spreading of expansion, while the more distal interactions of interior chromosomes are not tethered, and thus are lost during expansion. Alternatively, nucleolus association could influence the degree to which different chromosomes expand and spread during expansion.

**Figure 5.**
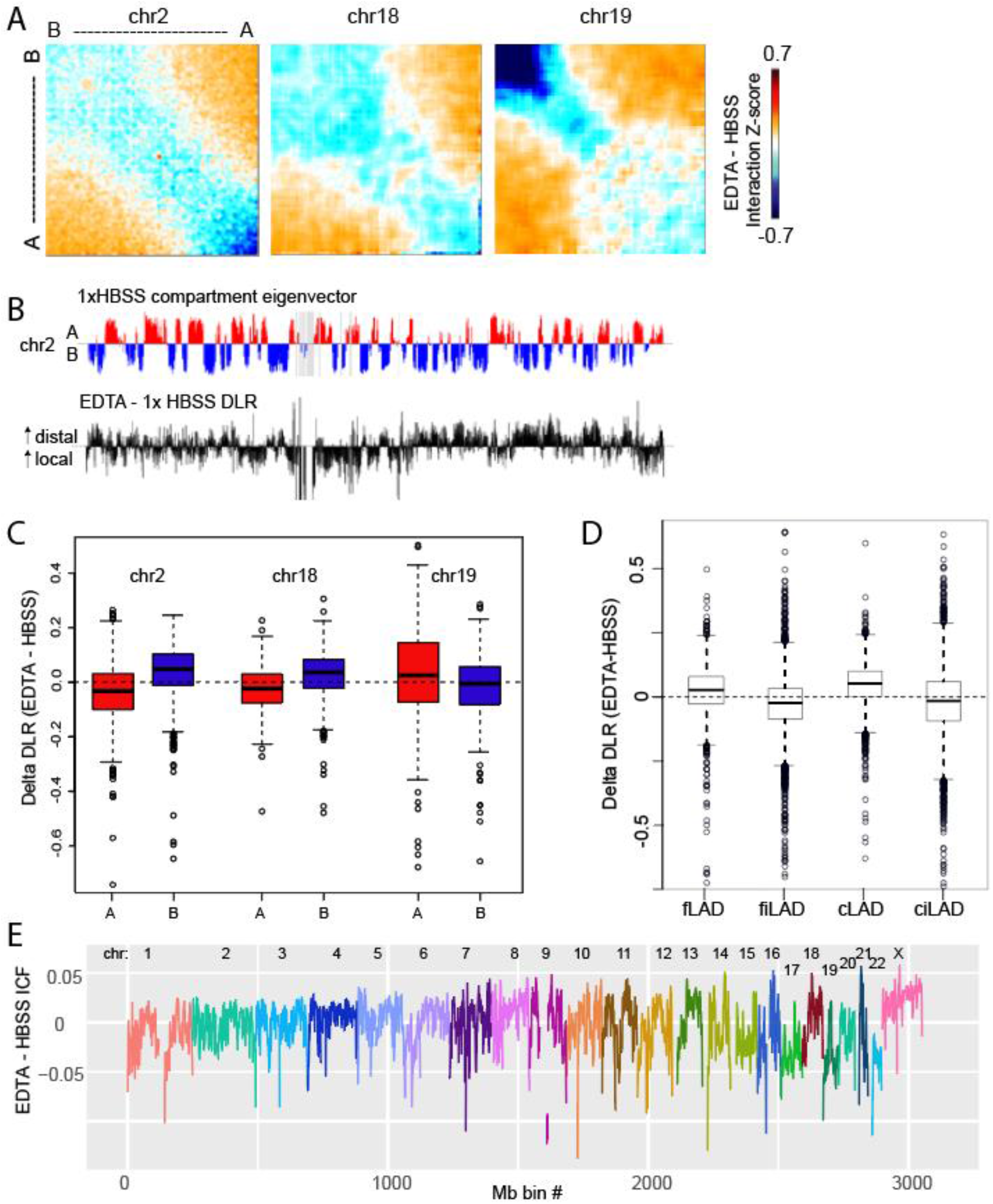
A) Saddleplots show interaction changes within and between compartments during EDTA expansion. Each matrix is sorted by compartment eigenvector value, from strongest B to strongest A compartment bins. Each pixel represents the difference between interaction Zscore EDTA - 1xHBSS. B) The distal to local ratio (DLR) was calculated as the sum of interactions greater than 3 Mb vs. less than 3 Mb for each 100 kb bin and then the difference in DLR was calculated between EDTA and HBSS and median centered. Regions of increased and decreased DLR visually correspond to B and A compartments. C) DLR differences segregate by compartment in LAD associated chr2 and chr18 but not NAD associated chr19. D) Changes in DLR for the whole genome segregated by LAD status as classified by Kind et al. E) Interchromosomal fraction (trans/cis interactions) difference between EDTA and HBSS is plotted in 1 Mb bins across the genome. Each chromosome is a different color.

### Effects of expansion on loop extrusion domains

Aggregating the Hi-C interaction signal across previously documented loops in GM12878 cells (Rao et al. 2014) shows that there is a decrease of loop strength on average during expansion (Figure 6A). However, we observe that not all loops behave uniformly during expansion. Loops bound by transcription-associated factors such as TBP or P300 do not decrease as much on average as CTCF loops (Figure 6B). These factors, which have previously been implicated in enhancer promoter looping (Bertolino and Singh 2002; Liu et al. 2011), are enriched at loops without CTCF motifs at their anchors. A scatterplot of loop strength relative to local background signal shows that some loops appear to increase in strength while others decrease during EDTA expansion, and then most loops return closer to their original strength upon return to 1xHBSS (Figure 6C). When we examined these differential changes in detail, we find several different classes of change that occur in different regions. We observe two categories of loops that decrease in contact frequency with expansion. In one case, we see strong interactions that decrease, while surrounding interaction levels are maintained (Figure 6D). In the other case, all interactions in a given region decrease uniformly, lessening the apparent loop, and also decreasing interactions within TADs (Figure 6E). We also observe a set of loops that increase in contact frequency with expansion, becoming more defined relative to the interactions within the TAD that this loop circumscribes (Figure 6F). In some of these cases, the loop strength is maintained while the “lines” surrounding the strong interaction are decreased (Figure 6G). These lines have been attributed to the effects of loop extrusion (Fudenberg et al. 2016), with higher resolution datasets revealing enriched sites of contact along these lines where loop extrusion “pauses” at intermediate CTCF sites and enhancer-promoter interactions (Krietenstein et al. 2020). The loss of lines and maintenance of “dots” may indicate that, upon the induced chromatin decompaction, cohesin rings slide until blocked by a CTCF site.

**Figure 6.**
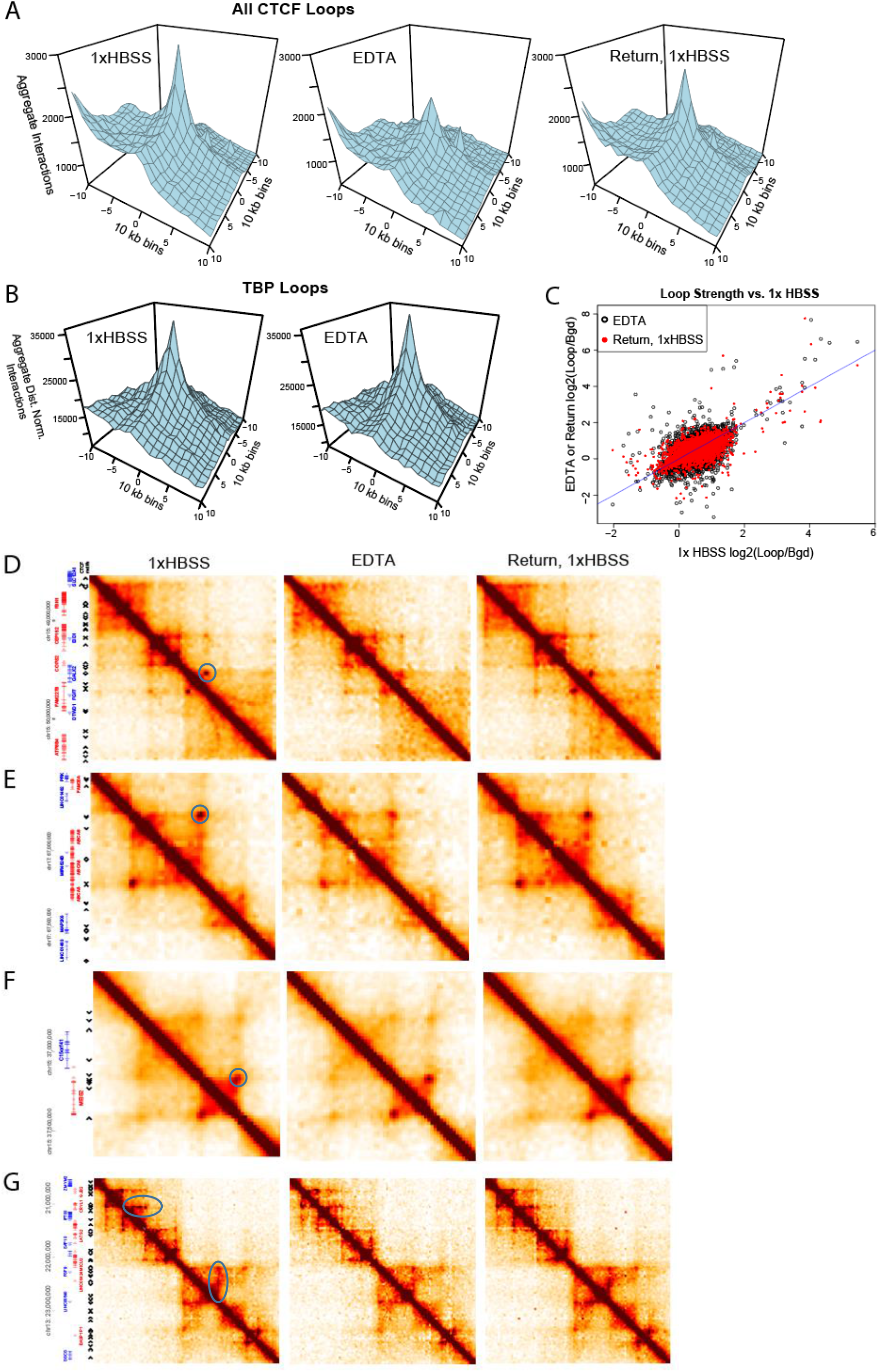
A) Aggregated interactions across all previously annotated loops (Rao et al., 2014) that contain CTCF sites at both anchors. B) Aggregated log2(observed/expected) plots for loops containing TBP binding at both loop anchors as measured by ENCODE in GM12878. C) Each point represents the log ratio of interactions in the 30 kb bin containing the loop vs. the interactions in a donut around this loop (bgd). Red points represent loop strength upon return to 1x HBSS vs. the original 1x HBSS strength while black points represent EDTA loop strength. D-G) Variations in loop strength during expansion in different regions. Each plot is binned at 20 kb, smoothed to 40 kb. Gene positions and CTCF motif positions shown at left. Areas of change highlighted in blue circles in 1xHBSS plot D-E) Decreased interactions. F) Increased interaction. G) Loss of loop extrusion lines during expansion.

The different classes of local interaction change we see do not segregate simply with compartment status or CTCF occupancy. Further investigation will be needed to clarify the different classes of local interaction change that we observe, but we hypothesize that in each case, the more highly reinforced interactions are being preserved during expansion, while contacts that are “transiently or incidentally proximal” are lost. Notably, each of these types of interaction changes are reversible upon return to 1x HBSS, even though the expanded state was maintained for an hour.

Our Hi-C data overall indicates that while many structural features are preserved during expansion, some contacts are lost. To further test this conclusion experimentally, we treated nuclei with increasing concentrations of formaldehyde before expansion. Progressively higher concentrations of formaldehyde rapidly inhibit expansion upon EDTA treatment (Figure S6A). This further suggests that some interactions which are present originally in 1xHBSS (and covalently linked by formaldehyde) must separate in order for full expansion in EDTA to occur.

### Nucleus expansion requires initial chromatin compaction and intact chromosome fibers

Our Hi-C results indicate that different chromatin types (A and B compartments and subcompartments) behave differently with expansion. To test the idea that initial chromatin modification state affects nucleus expansion, we treated either intact cells or isolated nuclei with 0.5 μM of the histone deacetylase inhibitor trichostatin A (TSA) for either 2 or 24 hours. This treatment has been widely used for its ability to increase histone acetylation and decompact chromatin (Wang et al. 2009; Bustos et al. 2017), and has been shown to make nuclei less stiff and more able to be stretched by external forces (Stephens et al. 2018). Even in isolated nuclei, real time chromatin modification changes have been seen upon treatment with histone deacetylase inhibitors (Sardo et al. 2017). In isolated nuclei, antibody staining for histone modifications can be performed without crosslinking (Sardo et al. 2017), allowing us to monitor the same stained nucleus across expansion conditions. TSA treated nuclei were already a bit increased in size in 1xHBSS as compared to controls, and then showed very little expansion upon buffer switch into EDTA (Figure 7A-C). This result agrees with the previously suggested idea of the prestressed state of the nucleus (Mazumder et al. 2010a), indicating that the chromatin fiber normally acts as a compressed spring, both resisting external forces and springing outward upon salt change. Once TSA treatment has already released this compressed spring energy by decondensing the chromatin, little further expansion is possible in low salt conditions. Interestingly, TSA treated nuclei never reach the size of low salt expanded nuclei (Figure 7B), suggesting that increased acetylation decondenses the genome in a way that allows more intermingling of chromatin domains, reducing the polymer persistence length, and thus the chromosomes exert less force on the nucleus exterior than low salt expansion with maintained chromosome topology.

**Figure 7.**
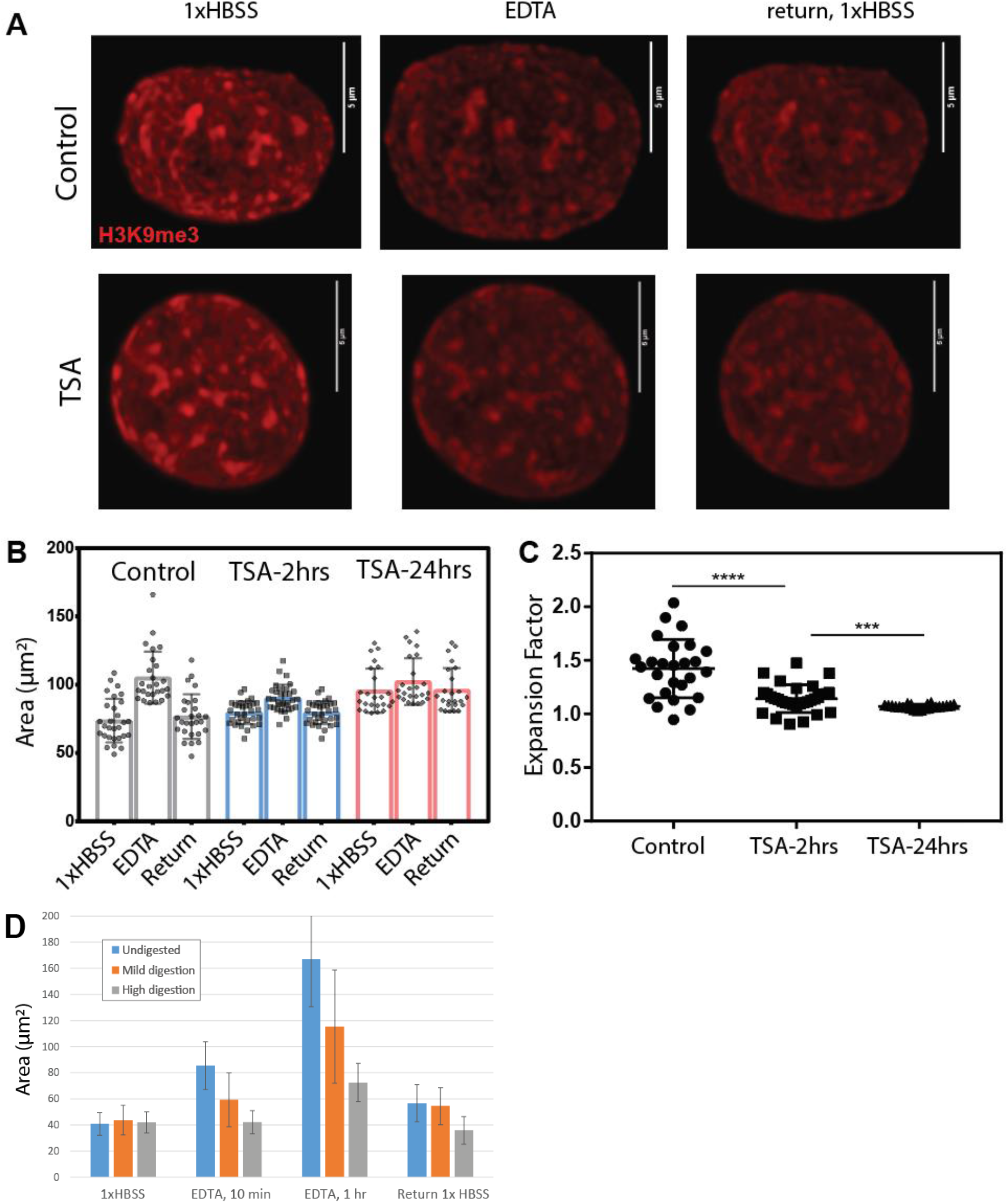
A) Isolated A375 cell nuclei were treated with TSA for 2 or 24 hrs, stained without crosslinking with H3K9me3, and then expanded with EDTA. Images show maximum projection of a confocal Z stack; scale = 5 µm. B) Area of each nucleus in each condition. Median +/-std dev with full range of data shown as points. C) Expansion factor = (EDTA / 1x HBSS nucleus area) for each nucleus analyzed. D) Isolated nuclei with no treatment, mild digestion (HindIII for 4 hr, 37 C) or high digestion (DpnII for 4 hr, 37 C) before expansion.

Also in line with the idea of the chromatin fiber as a compressed spring, we find that digesting DNA inside isolated nuclei with an endonuclease (Belaghzal et al. 2021) also inhibits nucleus low salt expansion (Figure 7D). Moderate digestion with a 6-cutter enzyme tempers the nucleus swelling somewhat, while digestion with a more frequently cutting 4-cutter enzyme effectively prevents nucleus expansion. Previous results show that this degree of 6-cutter digestion cuts the genome into 10-15 kb fragments, but that the genome structure is still held together by relatively many stable associations at this level of digestion (Belaghzal et al. 2021). Our expansion results suggest that with an intact fiber, or at least a sufficient degree of maintained associations (mild digestion), the expansion of the fiber causes the whole structure to be forced outward. However, when the chromosome fiber is further fragmented, local domains can expand and intermingle with each other and expansion does not occur.

### Imaging stained chromatin in expanded nuclei reveals additional chromatin structure detail

We observed that 3D genome contacts are largely preserved in the expanded nucleus, while average distances between genomic regions increase. Thus, we predicted that imaging expanded nuclei labeled for certain chromatin types or regions should provide visually more distinct chromatin structure information. Combining this idea with the fact that fluorescent immunostaining of histone modifications can be performed without fixation in isolated nuclei (Sardo et al. 2017), we stained isolated GM12878 nuclei for H3K9Ac, a mark of active chromatin. Before expansion, the staining pattern fills most of the nucleus in a diffuse and overlapping pattern. But, after 10 minutes of low salt expansion, distinct puncta are visible, possibly representing foci of active genes. The original staining pattern is restored upon re-contraction, confirming again that structures were not broken or rearranged during expansion (Figure S6B).

## Discussion

Overall, our results reveal both the overall robustness of the 3D genome structure to low salt nucleus swelling and demonstrate the utility of this expansion approach to reveal differences in the properties of different types of chromosome interactions. Nucleus expansion and re-contraction does not lead to major systematic rearrangements in compartment identity, chromosome positions, or TAD boundary positions. The reversibility of the expansion condition that is observed microscopically is borne out at the level of chromosome contacts as well. Interestingly, this result coincides with the computational predictions obtained from polymer physics models. Polymer models have predicted that local changes at the scale of unfolding the 30 nm fiber would not affect the larger scale conformations of the chromosome (Florescu et al. 2016). These results suggest that fundamental polymer properties of the genome allow local chromatin unfolding without dramatic disruption of larger scale chromosome organization.

We observe changes in the local decay of interactions with genomic distance upon expansion, and while these changes create a notable overall shift in TAD insulation profiles, TAD boundary positions are maintained. This suggests that genome topology can be largely independent of chromosome compaction. Indeed, genome topology may be tuned to facilitate robustness to nuclear shape change. The elastic expansion observed in normal nuclei can be destroyed either by strengthening connections through mild crosslinking or by removing connections by causing DNA breaks. This suggests that the genome topology maintains a level of connectivity that promotes both the elasticity and robustness of the resulting structure. Such a pre-loaded spring state may be key to the support of the physical structure of the nucleus by the 3D chromatin folding (Stephens et al. 2017b; Stephens et al. 2018). Strikingly, the weakening of connections, particularly B compartment interactions, that we do observe during EDTA induced expansion are similar to the loss of B compartment interactions observed after cells have deformed their nucleus to squeeze through tight constrictions (Jacobson et al. 2018; Golloshi et al. 2020). This suggests that there may be shared features of chromosome structure change during nucleus deformation in general.

These findings also have implications for biological circumstances in which visible expansion and contraction of nuclei are observed. Here, we have used changes in cation concentration as an artificial perturbation to examine chromosome structural properties, but such cation changes can also happen in vivo. Chromosome compaction upon ATP depletion and expansion upon return to normal ATP levels has been shown to depend on changes in cation and polyamine concentrations (Visvanathan et al. 2013; Kirmes et al. 2015). Often, when these large visible changes in the size of the nucleus are observed *in vivo*, it is assumed that the topology of genome contacts must be fundamentally rearranging. Our results would suggest that such changes might actually be accommodated without causing substantial rewiring of genome contacts. On the other hand, future work is needed to determine whether the subtle changes we observe in contact frequencies at certain loops and within TAD substructures could in fact have important biological outcomes, even while so much of the genome organization is unchanged. It is also important to note that our observations of the robustness of 3D genome contacts during low salt expansion are true on average across the population of nuclei. However, even when the mean structure does not change, there may be increased variation around the mean during low salt expansion across individual single cells. Future work with single cell Hi-C experiments and chromosome fluctuation analyses (Lindsay et al. 2018) will be necessary to clarify this point.

Beyond relevance to physiological circumstances, our results provide another tool to perturb the nucleus and observe physical aspects of the genome structure. These results can be incorporated into future models that aim to explain chromosome structure, such as the loop extrusion model of TAD formation. It is particularly striking that CTCF loops, even though sometimes weakened during expansion, are overall preserved or restored upon return to original salt conditions. Our results also provide a baseline for future work using expansion to perturb and assay effects on the chromosome structure. Future work will be needed to determine whether the subtle changes in structures we observe at 1 hour of expansion would become exacerbated by additional time in this expanded state. Starting from our observations of the overall robustness of genome structures to expansion, future work is also needed to define what architectural proteins and other factors are key to maintain this structure under expansion stress.

Finally, our results reveal subtle, but consistent differences in how different regions of the genome respond to expansion. We consistently observe different contact frequency changes between peripheral, lamin associated, gene poor chromosomes and internal, gene dense, nucleolus associated chromosomes. These two categories of chromosomes are differently affected by expansion in their local scaling changes, TAD insulation profile changes and inter-vs. intra-chromosomal interaction frequencies. More work will be needed to determine how much these differences are dependent on physical tethering to structures like the lamina and nucleolus and whether they relate to local histone modification differences. Pointing to the potential importance of local chromatin state, we observe that global histone acetylation increases decrease the propensity of the nucleus to expand at all in low salt. This result is in line with our observations that it is the B compartment (inherently lower histone acetylation to begin with) that decreases interactions more with expansion than the A compartment (which has a higher initial acetylation state). Further, we observe that different types of looping interactions respond differently to expansion. Future work is needed to clarify whether these different classes of loops are formed by different processes and what factors contribute to their different properties.

## Methods

### Cell lines

GM12878 cells were acquired under a Material Transfer Agreement from the Coriell Institute and were cultured in RPMI with 15% FBS. U2-OS, HEK293T, IMR90, B16-F1, and A375 cells were acquired from ATCC and cultured as recommended.

### Plasmid constructs

The Dendra2-Histone H4 plasmid was derived in part from a photoactivatable GFP-Histone H4 plasmid that was a kind gift of Dr. Roeland Dirks. The H4 gene was isolated from this plasmid by PCR using primers containing KpnI and BamHI sites:

5’ end of Histone H4 + KpnI: 5’-GCGCGCGGTACCATGTCTGGTAGAGGCAAAGG-3’

3’ end of Histone H4 + BamHI: 5’-GCGGATCCCGGGTCAGCCACCAAAGCCGTACA-3’.

Histone H4 was then cloned into the multiple cloning site of the pDendra2-C vector (Clontech catalog# 632546). The dCas9-3xGFP and C9-sgRNA plasmid for CRISPR imaging were a kind gift of Dr. Hanhui Ma and Dr. Thoru Pederson. The GM12878 cells were transfected with pDendra2-H4 using Kit L and program Y-001 on a Lonza Nucleofector IIb. B16-F1 cells were transfected with pDendra2-H4 using the Attractene Fast-Forward protocol (Qiagen). U2-OS and HEK293T cells were transfected with dCas9-3xGFP and C9-sgRNA plasmid at a 1:10 ratio using Lipofectamine 3000 (Thermo Fisher).

To carry out expansion experiments on progerin expressing nuclei, IMR90 cells were transfected with 1 µg of pEGFP-Δ50 LMNA (Addgene Plasmid #17653). After expression of the GFP was observed (∼2 days), nuclei were isolated and expansion experiments carried out as described below.

### Nuclei Isolation

GM12878 nuclei were isolated with the following approach. 1×10^8^ cells were collected by centrifugation (200xg, 5 minutes, room temperature) and then washed twice with 10 mL ice-cold 1X dPBS and then twice with 5 mL ice cold Nuclei Buffer (NB) (10 mM PIPES pH 7.4, 2 mM MgCl_2_, 10 mM KCl, 1mM DTT, H_2_O pH 7.4) containing protease inhibitors (Thermo Scientific). After final wash and spin, the cell pellet was resuspended in 10 mL NB + 0.06% NP-40, vortexed gently, and then allowed to equilibrate on ice for 10 minutes. Cells were then homogenized on ice using a Dounce homogenizer (Pestle A) with 10 slow strokes, rest for 20 minutes on ice, then 10 more strokes if needed. The cells were examined under a microscope after each 10 strokes to evaluate the degree of lysis and nucleus release. Cells may lyse with 10-30 strokes. To remove excess cellular material, 5 mL of lysed cell mixture was layered on top of a 20 mL sucrose cushion: 30% sucrose in NB and centrifuged for 10 minutes at 730xg. The supernatant and interface was carefully removed and the pellet containing nuclei was resuspended in either 1x Hank’s Balanced Salt Solution (HBSS: 1.26 mM CaCl2, 0.49 mM MgCl2, 0.41 mM MgSO4, 5 mM KCl, 0.44 mM KH2PO4, 4.17 mM NaHCO3, 137.9 mM NaCl, 0.34 mM Na2HPO4, 5.56 mM D-Glucose) for immediate use, or in Nuclei Storage Buffer pH 7.4 (10 mM PIPES pH 7.4, 20 mM NaCl, 80 mM KCl, 50% glycerol, 250 mM sucrose, 0.5 mM DTT, 1x protease inhibitors) for storage at - 80°C for later use.

U2-OS, HEK293T, A375, IMR90, and B16-F1 nuclei were isolated using the same method as previously described after detaching cells with trypsin. Adherent cell types typically required more Dounce homogenization strokes than suspension cells.

### Expansion and Contraction Conditions

Isolated nuclei in 1x HBSS were plated onto 35 mm poly-D-lysine coated glass bottom dishes (MatTek P35GC-1.5-10-C) for imaging or poly-D-lysine coated 100 mm culture plates for Hi-C experiments (Corning BioCoat 356469), and maintained at 4° C for a few hours or overnight until nuclei were settled and adhering to the dish. Nuclei were expanded by removing 1xHBSS carefully with a pipette until only a thin layer of liquid remained (complete drying would destroy nuclei) and washing nuclei with 1 mM EDTA in 10 mM HEPES buffer pH 7.4 three times. Imaging occurred immediately upon addition of EDTA and in time intervals ranging from 10 minutes to an hour. In one set of experiments (including pDendra labeled nucleus expansion in Figure 1), 25 µg RNAse in 200 uL 1x HBSS was added to nuclei for 10 minutes before EDTA expansion was carried out. We found no detectable effect of this RNAse treatment either on the photoconverted pattern or the expansion properties. All other experiments shown were performed without RNAse.

For expansion experiments on cells pre-treated with TSA, A375 cells were treated with 0.5 µM TSA for either 2 hours or 24 hours before nuclei isolation. Then, isolated nuclei were settled into the PDL coated dish and expansion was carried out as usual (1xHBSS, then 10 min EDTA and then 10 min recovery in 1xHBSS).

Hi-C experiments were performed after 1 hour of EDTA expansion. Nuclei were returned to their contracted pre-EDTA state by removing EDTA and washing with 1x HBSS three times. The contracted state was then imaged immediately afterward and in time intervals ranging from 10 minutes to an hour. Hi-C experiments were performed 20 min after return to 1x HBSS. One Hi-C replicate was performed with expansion performed as above, but replacing 1x HBSS with 0.1x HBSS instead of HEPES-EDTA.

### Antibodies and isolated nucleus staining

To stain isolated nuclei for heterochromatin marks, nuclei were settled in a PDL coated dish in 1xHBSS and then stained with a protocol adapted from (Sardo et al. 2017). Specifically, nuclei were incubated for 1 hour in a solution containing 5% FBS and 1:500 primary antibody (H3K9me3; Invitrogen rabbit polyclonal 49-1008 or H3K9Ac; Invitrogen 49-1009) in 1x PBS. The nuclei were then washed twice with PBS 5% FBS and then incubated with secondary antibodies (Fisher Goat anti-Rabbit Secondary Antibody Alexa 488 or Alexa Fluor 594; R37116 or R37117) in PBS 5% FBS for 1 hour. Finally, the nuclei were washed twice with PBS 5% FBS. Stained nuclei could then be treated with EDTA for expansion.

### Imaging and Image Analysis

Changes in nucleus size with expansion were quantified by taking images at 40x magnification on an EVOS FL (Thermo Fisher) microscope. Expansion was typically performed on the microscope stage so that the same nuclei could be captured before, during, and after expansion. Nucleus dimensions were quantified using ImageJ by manually fitting an ellipse around each nucleus.

Patterns were drawn on pDendra2-H4 expressing GM12878 nuclei typically using 6-10 scans of 4-5% power of a 405 nm laser on a Leica Confocal SP8 microscope. CRISPR-GFP images were collected at the same intensity of a 488 nm laser for all conditions, and a stack of images from the top to the bottom of each nucleus was acquired. The intensity of fluorescent foci was quantified in ImageJ using the maximum projection image. The average of 3 background measurements were subtracted from the mean fluorescence across each spot area. All measurements were expressed as a fraction of the initial intensity of the spot in the 1xHBSS condition and then normalized by the average intensity loss of spots that were not expanded, but only imaged 3 times in the 1xHBSS buffer (to account for the mild photobleaching after taking multiple stacks of images).

### mFISH

mFISH using human chromosome-specific probes were essentially the same as described by the manufacturer (MetaSystems,70 Bridge St., Suite 100, Newton, MA 02458 USA) and in previous studies (Balajee et al. 2018). Briefly, slides were treated for 1 min with 0.001% acidic pepsin solution (0.01 N HCl) at 37°C for 1–2 min followed by two washes of 5 min each in PBS. The slides were fixed for 10 min in a solution of formaldehyde/MgCl2 (1% formaldehyde/50 mM MgCl2in PBS). The slides after denaturation were dehydrated in graded series of ethanol (30%, 70%, 90%, and 100%) and air dried. The mFISH probes was denatured separately by incubation at 75°C for 5 min followed by incubation at 37°C for 30 min to allow the annealing of repetitive DNA sequences. An aliquot of the 10mL probe was placed on the slide and covered with a coverslip. The slides were kept in a humidified hybridization chamber at 37°C for at least 72 h. The unbound probe was removed by washing the slides in prewarmed (75°C) SSC (pH 7.0–7.5) for 5 min followed by incubation in 4xSSCT (4xSSC with0.1% Tween 20) for 5 min. Images of interphase nuclei and metaphase chromosomes were acquired using the Zeiss epifluorescence microscope equipped with a 63x objective and five optical filters for the excitation of five different fluorochromes [FITC, Spectrum Orange, Texas Red, DEAC, and Cy5]. Territories specific for each homologous pair of chromosomes were identified by unique chromosome-specific processed color generated by the Isis software (MetaSystems). For acquisition, a single two-dimensional image was captured using multiple filters.

### Digestion and Crosslinking before Expansion

To test the effect of intact vs. fragmented chromosomes on expansion, isolated nuclei were, subjected to either 50 U HindIII per µg of DNA (mild digestion) or 10 U DpnII per µg of DNA (high digestion). Nuclei were allowed to settle on a PDL coated coverslip dish and then the density of nuclei per area was estimated under a 40x objective. The amount of DNA in each coverslip dish was estimated as (# nuclei/mm^2^) × (314 mm^2^/dish) × (6.2× 10^−6^ µg DNA/nucleus). This DNA quantity was used to calculate the amount of restriction enzyme to use. Nuclei were incubated with restriction enzyme in 1x CutSmart buffer (New England Biolabs) for 4 hr at 37 °C and then washed with 1xHBSS before expansion.

To test the effect of formaldehyde crosslinking on nucleus expansion, nuclei were settled in PDL coated coverslip dishes and then crosslinked with either 0.25%, 0.5%, 1%, 2%, or 5% formaldehyde in 1x HBSS for 10 minutes at room temperature. Crosslinking was quenched with glycine and then dishes were washed with 1xHBSS before expansion experiments.

### Hi-C

Hi-C experiments were performed essentially as described previously, but with a few important modifications. 10-20 million cells were used for each condition. Crosslinking was performed with 1% formaldehyde pre-mixed with the corresponding appropriate buffer (1x HBSS, or 1 mM EDTA + 10 mM HEPES) immediately before addition to the plates of nuclei. After 10 minutes, crosslinking was quenched with glycine, and plates of crosslinked nuclei were washed in the same buffer corresponding to their expansion or contraction condition. It was not possible to harvest and collect nuclei by scraping and centrifugation due to the fragility of the nuclei, particularly in the expanded case, so nuclei were lysed directly in plates by aspirating all liquid and then adding 1.8 mL of 0.3% SDS in 1x NEB2 buffer (New England Biolabs) per 10 cm dish. Dishes were incubated with gentle agitation at 37°C for 30 min. Crosslinked material was then removed the plate by scraping. If more than one plate was to be combined to obtain enough cells, the SDS-cell mixture scraped from the first plate was transferred to the second plate for another round of incubation, so that cells from all plates were combined into a final volume of less than 3 mL. SDS was sequestered by addition of 1.8% Triton (final concentration). Crosslinked complexes were then divided into aliquots of 560 uL each to match other Hi-C protocols, and were then subjected to digestion with HindIII overnight (500 U per 560 uL aliquot), biotin fill-in, and dilute ligation as previously described (Belton et al. 2012).

### Hi-C Sequencing and Contact Matrix Construction

Hi-C libraries were sequenced on an Illumina HiSeq platform with paired end reads. Sequencing reads were mapped to the human genome (hg19), filtered, and iteratively corrected as previously described (Imakaev et al. 2012) (https://github.com/dekkerlab/cMapping). For library quality and mapping statistics see Table S1.

### Hi-C reproducibility assessment

A spearman correlation-based metric was used to assess reproducibility across all Hi-C datasets, as previously described in Sanders et al. (Sanders et al. 2020). After taking the log of iteratively corrected 1 Mb binned interaction counts and excluding the diagonal bin, correlation maps were calculated for each individual matrix. Each entry in such a Hi-C data correlation matrix represents the correlation between the full row and column of data at that position in the matrix across the entire genome. These individual condition correlation matrices were then used as input to the calculation of pairwise correlations between datasets.

### Interchromosomal Interaction Analysis

The relative frequencies of interactions between each chromosome was determined by comparing the observed number of Hi-C reads involving interactions between two chromosomes to the number of interactions expected to occur between them according to the total numbers of reads observed on each chromosome. This calculation was performed using the equation previously described (Zhang et al. 2012) using the collapseMatrix.pl script available at https://github.com/dekkerlab/cworld-dekker.

### Compartment analysis and compartment strength saddle plots

Compartment analysis was performed by running principal component analysis using the matrix2compartment.pl script from https://github.com/dekkerlab/cworld-dekker. Since the PC1 sign (positive or negative) is arbitrary, A compartment identity was assigned to whichever group of bins had the higher gene density. The PC1 value was then used to determine compartment identity for 100 and 250Kb binned matrices. Saddle plots were constructed to investigate changes in interaction frequency between or within compartments Interaction zScores, normalizing for interaction decay with genomic distance, were calculated according to the matrix2compartment.pl at 250 kb bin size. Then Zscore matrices were reordered based on compartment strength (from strongest B to strongest A using PC1 values). The control condition was subtracted from the expanded condition and then the final matrices were smoothed at a 500Kb bin size.

### Insulation Score and TAD Analysis

Topologically associating domain (TAD) boundaries were called using the InsulationScore method (Crane et al. 2015), available as matrix2insulation in the cworld-dekker github package. The insulation score was calculated at two different resolutions: 20 kb matrix bin size with a 160 kb insulation square size (the distance within which interactions would be summed) and 40 kb bin size with a 520 kb insulation square size. The matrix2insulation script then calls TAD boundaries as minima in the insulation score profile.

To calculate the average insulation score profile around sets of TAD boundaries, the computeMatrix and plotProfile tools were used from deepTools2 (Ramírez et al. 2016) as implemented in Galaxy Version 3.3.2.0.0 https://usegalaxy.org/. Sets of boundaries that occur in the A or B compartment were identified with the bedtools intersect function (Quinlan and Hall 2010). Subcompartment classifications of regions were used as previously calculated for GM12878 cells, downloaded from https://www.ncbi.nlm.nih.gov/geo/query/acc.cgi?acc=GSM1551550.(Rao et al. 2014)

### Interaction Scaling with Distance Analysis

Using 40 and 10 kb binned iteratively corrected and scaled contact matrices, we extracted contact frequencies between bins at each genomic distance, excluding the diagonal bin (zero distance). A loess fit was then used to find a smooth curve describing interaction decay vs. distance (using the matrix2loess.pl script in cworld-dekker). The interaction frequencies were then normalized to set the maximum value (loess fit interaction value for the minimum distance) for each dataset to 1 and then plotted on a log scale vs. log genomic distance. For 10 kb bin size, the maximum distance considered between two loci was capped at 1 Mb.

### Distal Local Ratio and Interchromosomal fraction

The ratio of interchromosomal interactions relative to the total number of interactions for a given chromosome was calculated as previously described (Heinz et al. 2018)

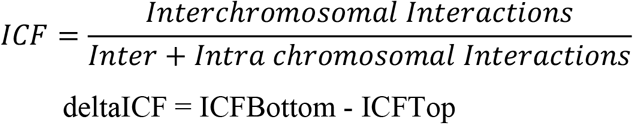

The log2 ratio of Hi-C interactions with distance greater than 3 Mb relative to local interactions less than 3 Mb away was defined as the Distal to Local Ratio. To find the change in DLR after expansion, delta DLR was calculated as DLR_EDTA_ – DLR_HBSS_ as previously described (Heinz et al. 2018).

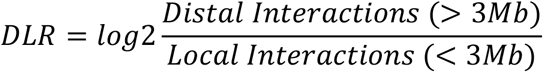

### Looping Interaction Analysis

For analysis of changes in looping interactions, we began with the set of loops previously defined from deeply sequenced GM12878 Hi-C data (Rao et al. 2014). We then classified these loops as CTCF containing loops if the loop anchor sites contained CTCF motifs on both sides according to the information in the Rao et al. supplementary table.

TBP site interactions were defined as follows. First, genomic locations binding TBP in GM12878 cells were assigned according to ENCODE data set wgEncodeAwgTfbsSydhGm12878TbpIggmusUniPk.narrowPeak (Consortium 2012) made available in the UCSC Genome Table Browser (http://genome.ucsc.edu/cgi-bin/hgTables) for hg19. Then, all possible combinations of Tbp binding sites separated by more than 50 kb were considered as potential looping interactions.

For a given set of interactions, the aggregate interactions were calculated as follows: Hi-C data, binned and corrected at 10 kb resolution for each condition and the corresponding looping interaction list was input into the interactionPileUp.pl script from cworld-dekker. This script extracts interaction sub-matrices 100 kb upstream and downstream of each loop anchor pair and then sums all interactions across all sub-matrices. We then further summed pile up matrices across all chromosomes to produce the final aggregated interaction plots.

To define the strength of each loop for plotting and comparing between conditions, we again used the set of loops defined by Rao et al and 10 kb binned Hi-C matrices for each condition. At each loop position, we calculated the ratio of the interactions in the loop bin divided by the sum of interactions of the bins surrounding the loop.

## Data Access

Hi-C data are available on GEO, accession number XXX

## Acknowledgements

We thank Sarah Carver, Trevor Freeman, and Jeff Schoondyke for assistance with some expansion experiments and quantification, and Houda Belaghzal for advice on digestion conditions. We thank Justin Lindsay for assistance with computational analysis, Bryan Lajoie, Hakan Ozadam, and Junwen Li for assistance with computational pipelines, and Noam Kaplan for helpful conversations. We thank Adayabalam Balajee for assistance with mFISH chromosome territory imaging. This work was supported in part by an NIH NIGMS grants F32GM100617 and R35GM133557 to R.P.M., and NIH grant HG003143 to J.D. J.D. is an investigator of the Howard Hughes Medical Institute.

## Author Contributions

JTS and RG performed Hi-C and imaging experiments, analyzed data, and contributed to writing the manuscript. PHT performed and analyzed digestion and CRISPR imaging experiments and contributed to writing the manuscript. DN performed and analyzed digestion expansion experiments. YX performed some data analysis. JD provided advice for designing the study and performing experiments, analyses, and writing the manuscript. RPM conceived the study, performed experiments, analyzed data, and wrote the manuscript.

## Supplementary Figures

**Figure S1**. A) DAPI staining after 1 h incubation of GM12878 nuclei in HEPES + 1mM EDTA (right) and return to 1x HBSS (right) shows chromatin fills expanded nucleus space (scale = 100 microns). B) The addition of DAPI after expansion reduces the size of the nuclei (EDTA 1h N = 115; DAPI N = 28; error bars = std. dev.). C) Photoconverted pattern is also preserved upon a different approach to low salt expansion (dilution to 0.1x HBSS) and in a different cell type (mouse melanoma; scale = 3 microns). D) Photoconverted pattern in GM12878 cell is preserved after 1 hour of expansion, though the nucleus rotated ∼90 degrees around the coverslip attachment point (see altered nucleolus position). (scale = 3 microns) E) RNase treatment does not affect nucleus expansion or fidelity of chromatin positioning after expansion. Nuclei were treated with 25 mg of RNAse (in 200 uL) for 10 min and then subjected to expansion. Scale bar = 2 µm.

**Figure S2**. A) IMR90 nuclei stably transfected with Progerin-GFP show wrinkles typical of progeria, but are able to expand and re-contract. B) Quantitative comparisons between WT LMNA-GFP and Progerin-GFP expressing IMR90 nuclei show a slight increase in initial size with progerin expression, but equivalent return to original size after expansion in both.

**Figure S3**. A) HEK293T cells transfected with Cas9-GFP targeting chr9 pericentromeric region. Spot distances and intensities for Figure 1F and G were quantified from these and similar images. B) No fluorescence recovery is observed after return to 1xHBSS when a Cas9-GFP spot is photobleached (white circle) in the expanded state. (scalebar = 3 microns) C) Expanded nucleus size is preserved after crosslinking with 1% formaldehyde. (Bars show mean, error bars = std. dev. N = 83, 71, 116, 28)

**Figure S4**. A) Genome wide Hi-C contact correlation at 1 Mb resolution between all datasets. Spearman correlation-based reproducibility is calculated as described in the Methods. Replicates have some systematic differences, but expanded conditions (EDTA and 0.1xHBSS) are the more separate from non-expanded conditions in each replicate set. B) Compartment identity is conserved for EDTA expansion replicate 2. chr2 eigenvector 1 is shown at 250 kb resolution. C) TAD insulation scores are preserved in expanded nuclei in both 0.1xHBSS and EDTA treatment replicate. Insulation scores shown for chr2 calculated with 520 kb insulation square size. D) Log10 interactions vs. Log10 of genomic distance in base pairs for all 40 kb binned intrachromosomal interactions genome wide in the 0.1xHBSS expansion condition and the second EDTA replicate. A faster drop in local interactions and more long range interactions is observed in expansion. E) Changes in Distal Local Ratio for the whole genome separated into LAD status as classified by Kind et al. As in Figure 5D, but using Hi-C data from EDTA replicate 2.

**Figure S5**. Average insulation profile (20 kb bin, 160kb insulation square) around TAD boundaries in either all B compartment regions (top) or in specific sub-compartments (defined in GM12878 cells by Rao et al., 2014).

**Figure S6** A) Nuclei were crosslinked with increasing percentages of formaldehyde for 10 min prior to expansion with EDTA and the average fold change in nucleus volume after EDTA for 10 minutes is indicated. B) H3K9Ac immunostained (green) isolated GM12878 nuclei show more detailed pattern of this histone modification localization in the EDTA expanded state. After return to 1xHBSS, the original pattern is largely recapitulated. Image shows a single confocal Z slice. Scale bar = 2 µm.

## Supplementary Tables

**Supp Table 1**. Hi-C read numbers and mapping/processing statistics

