## Supplementary Figures for "Loops, TADs, Compartments, and Territories are Elastic and Robust to Dramatic Nuclear Volume Swelling"

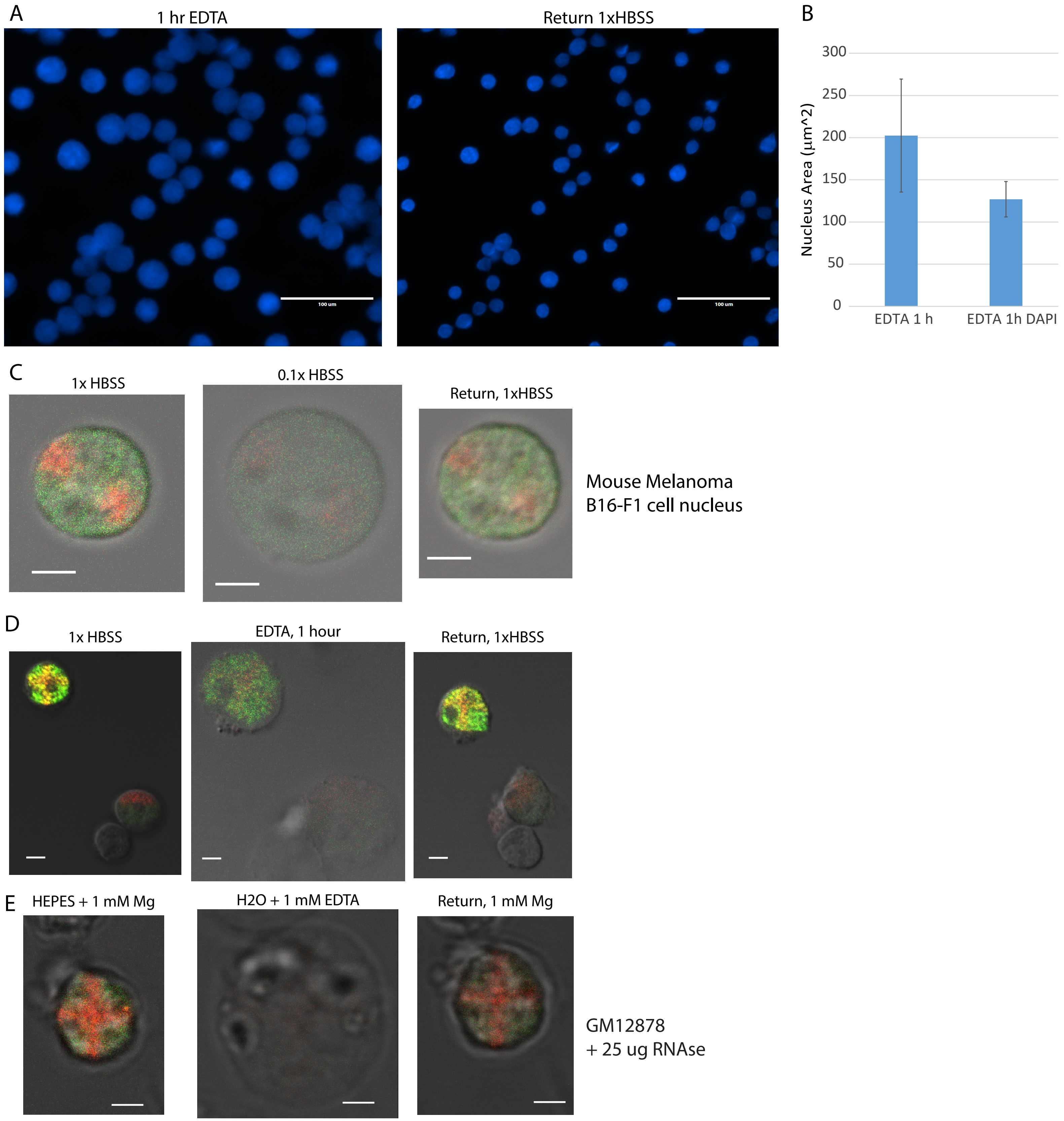


Figure S1. A) DAPI staining after 1 h incubation of GM12878 nuclei in HEPES + 1mM EDTA (right) and return to 1x HBSS (right) shows chromatin fills expanded nucleus space (scale = 100 microns). B) The addition of DAPI after expansion reduces the size of the nuclei (EDTA 1h N = 115; DAPI N = 28; error bars = std. dev.). C) Photoconverted pattern is also preserved upon a different approach to low salt expansion (dilution to 0.1x HBSS) and in a different cell type (mouse melanoma; scale = 3 microns). D) Photoconverted pattern in GM12878 cell is preserved after 1 hour of expansion, though the nucleus rotated ~90 degrees around the coverslip attachment point (see altered nucleolus position). (scale = 3 microns) E) RNase treatment does not affect nucleus expansion or fidelity of chromatin positioning after expansion. Nuclei were treated with 25 μg of RNAse (in 200 uL) for 10 min and then subjected to expansion. Scale bar = 2 μm.

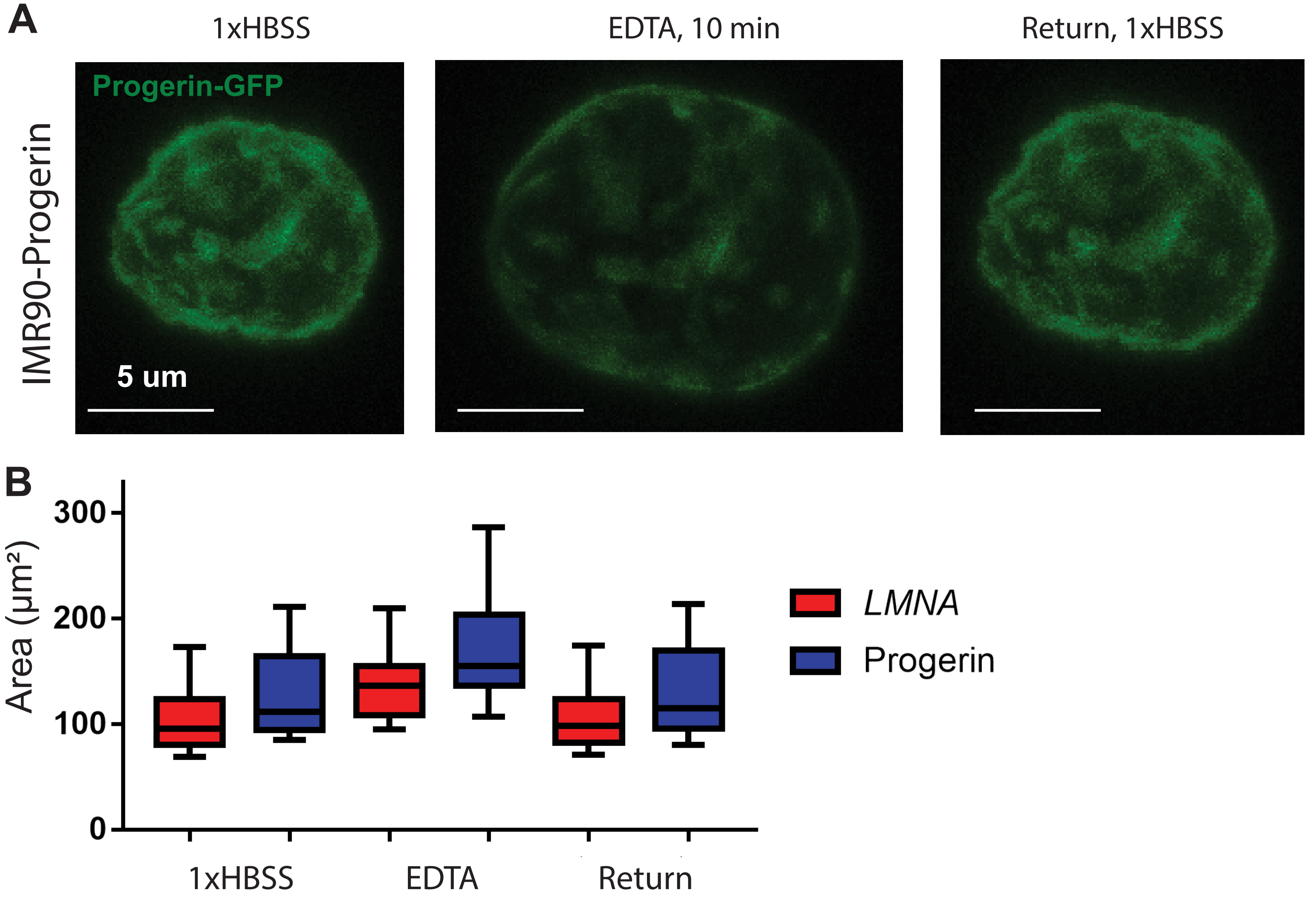


Figure S2 A) IMR90 nuclei stably transfected with Progerin-GFP show wrinkles typical of progeria, but are able to expand and re-contract. B) Quantitative comparisons between WT LMNA-GFP and Progerin-GFP expressing IMR90 nuclei show a slight increase in initial size with progerin expression, but equivalent return to original size after expansion in both.

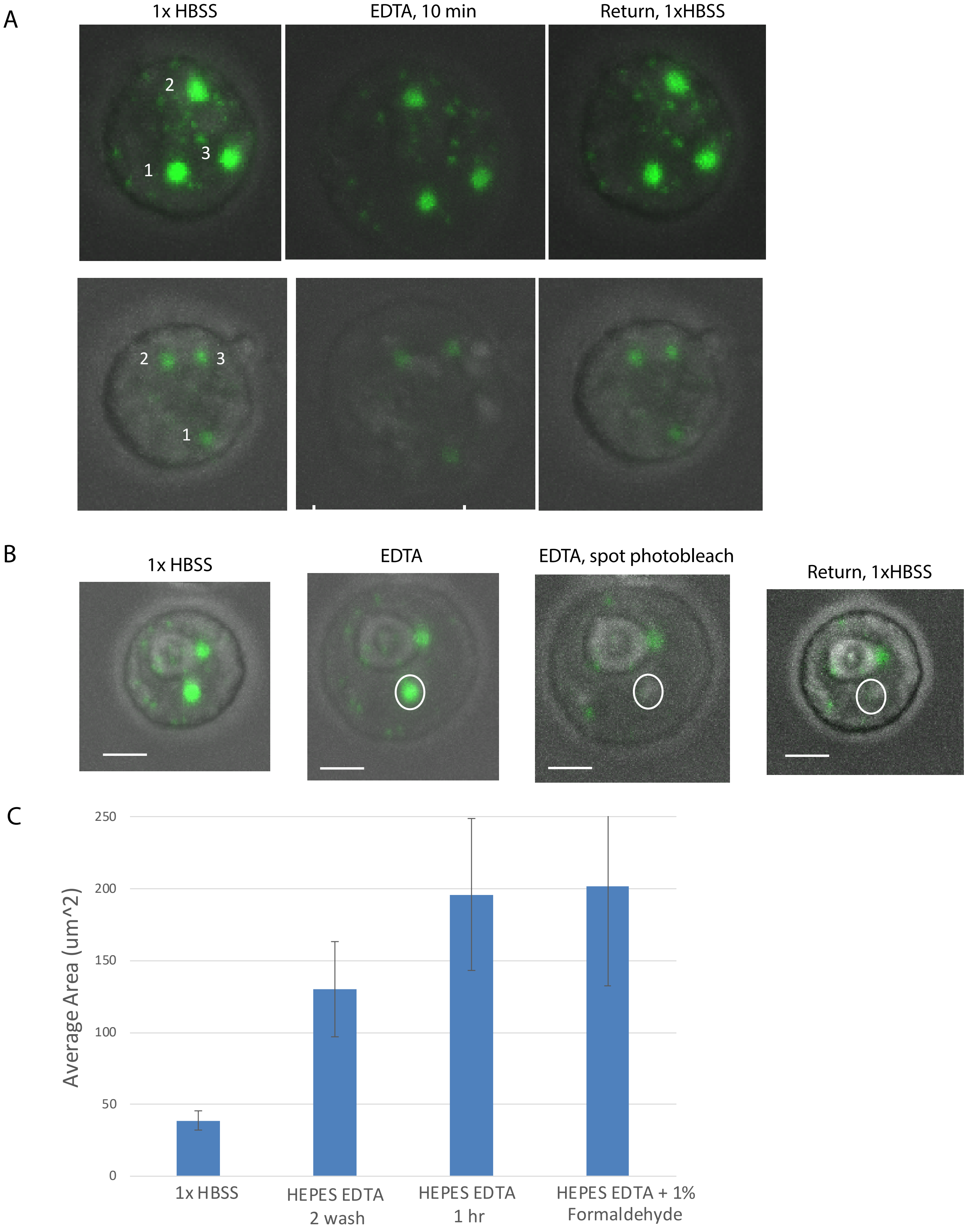


Figure S3. A) HEK293T cells transfected with Cas9-GFP targeting chr9 pericentromeric region. Spot distances and intensities for Figure 1F and G were quantified from these and similar images. B) No fluorescence recovery is observed after return to 1xHBSS when a Cas9-GFP spot is photobleached (white circle) in the expanded state. (scalebar = 3 microns) C) Expanded nucleus size is preserved after crosslinking with 1% formaldehyde. (Bars show mean, error bars = std. dev. N = 83, 71, 116, 28)

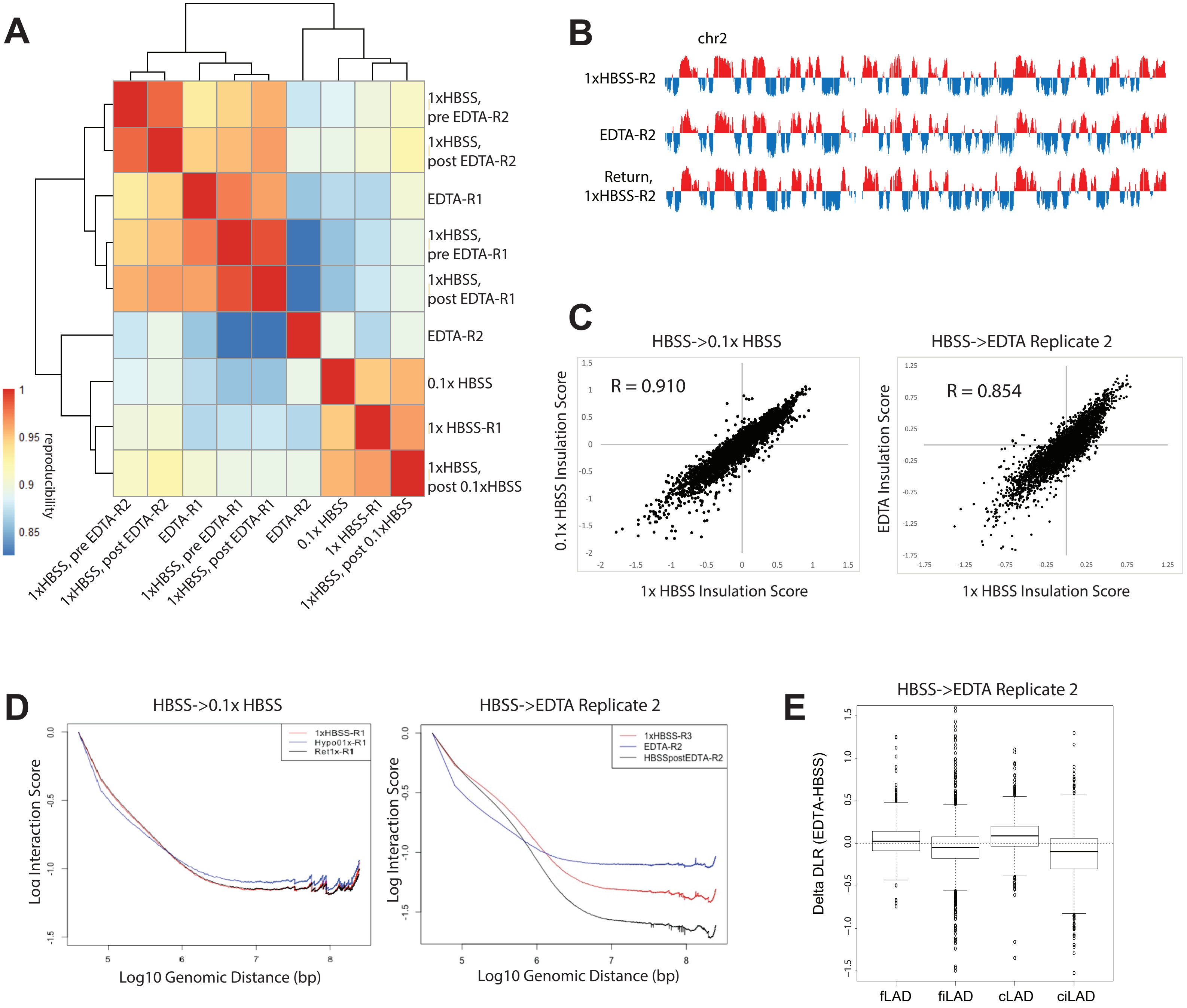


Figure S4. A) Genome wide Hi-C contact corelation at 1 Mb resolution between all datasets. Spearman correlation-based reproducibility is calculated as described in the Methods. Replicates have some systematic differences, but expanded conditions (EDTA and 0.1xHBSS) are the more separate from non expanded condtitions in each replicate set. B) Compartment identity is conserved for EDTA expansion replicate 2. chr2 eigenvector 1 is shown at 250 kb resolution. C) TAD insulation scores are preserved in expanded nuclei in both 0.1xHBSS and EDTA treatment replicate. Insulation scores shown for chr2 calculated with 520 kb insulation square size. D) Log10 interactions vs. Log10 of genomic distance in base pairs for all 40 kb binned intrachromosomal interactions genome wide in the 0.1xHBSS expansion condition and the second EDTA replicate. A faster drop in local interactions and more long range interactions is observed in expansion. E) Changes in Distal Local Ratio for the whole genome separated into LAD status as classified by Kind et al. As in Figure 5D, but using Hi-C data from EDTA replicate 2.

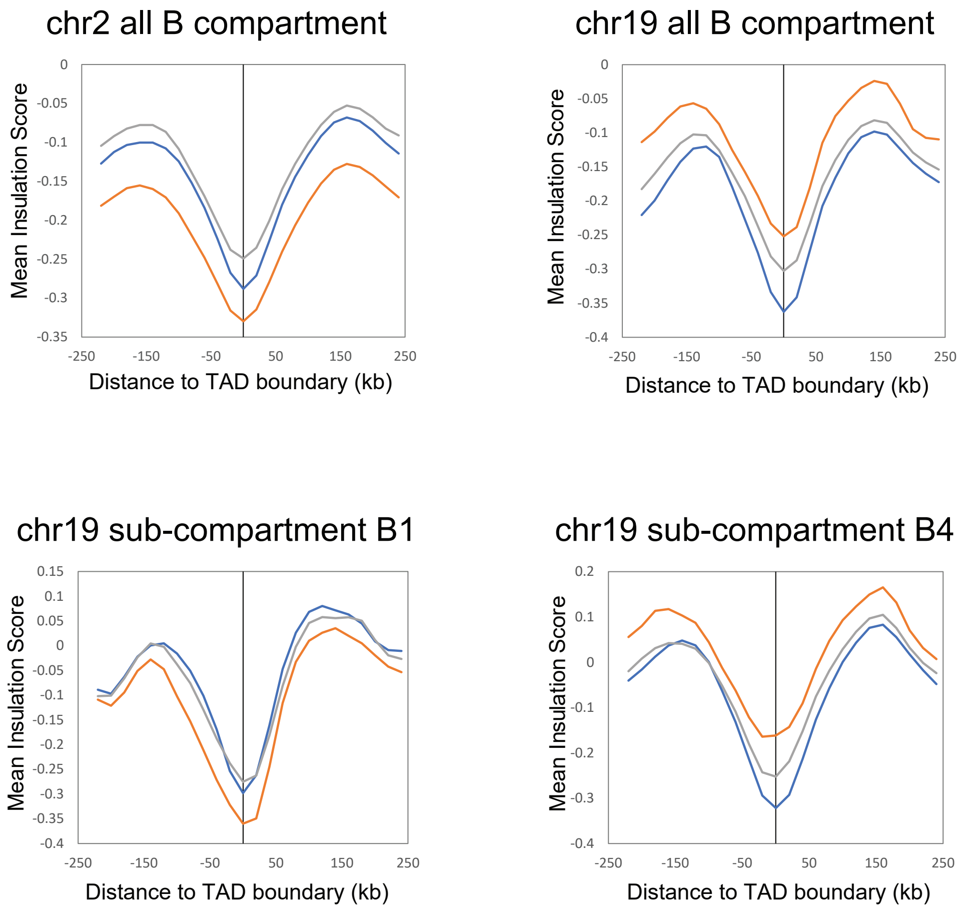


Figure S5. Average insulation profile (20 kb bin, 160kb insulation square) around TAD boundaries in either all B compartment regions (top) or in specific sub-compartments (defined in GM12878 cells by Rao et al., 2014).

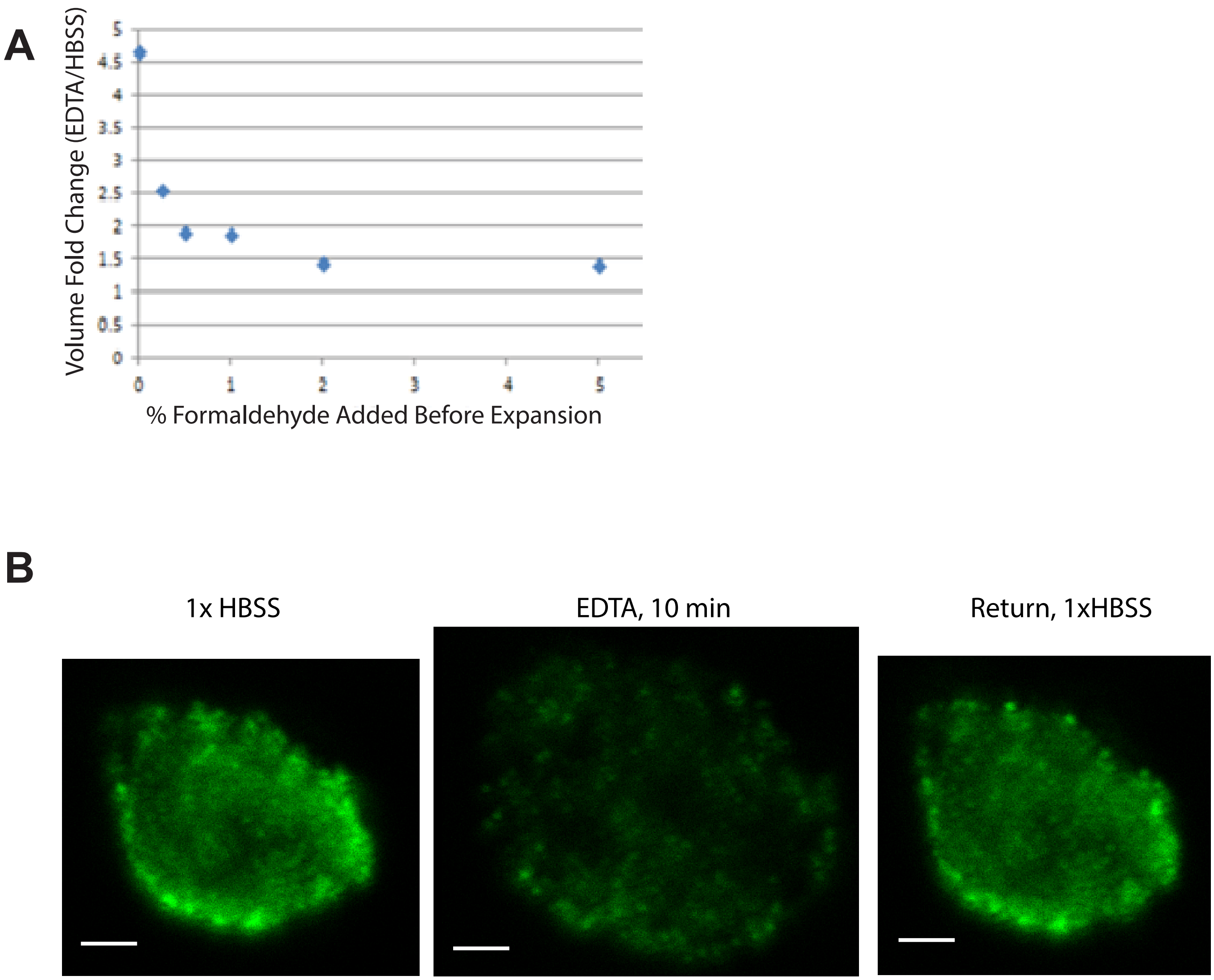


Figure S6 A) Nuclei were crosslinked with increasing percentages of formaldehyde for 10 min prior to expansion with EDTA and the average fold change in nucleus volume after EDTA for 10 minutes is indicated. B) H3K9Ac immunostained (green) isolated GM12878 nuclei show more detailed pattern of this histone modification localization in the EDTA expanded state. After return to 1xHBSS, the original pattern is largely recapitulated. Image shows a single confocal Z slice. Scale bar = 2 μm.
