## Supplementary Table 1 for "Loops, TADs, Compartments, and Territories are Elastic and Robust to Dramatic Nuclear Volume Swelling"

**Table S1. Hi-C Mapping Statistics**

| Condition | Replicate | Raw # Reads | # Both Sides Mapped | Valid Pairs | Unique Valid Pairs | % Cis | % Dangling Ends |
|---|---|---|---|---|---|---|---|
| 1x HBSS | R1 | 228,519,373 | 158,543,659 | 133,626,690 | 127,083,536 | 38.0 | 10.6 |
| 0.1x HBSS, 1 h | R1 | 226,837,794 | 162,358,348 | 117,359,572 | 88,661,306 | 44.3 | 22.4 |
| Return, 1xHBSS | R1 | 235,448,026 | 162,146,341 | 137,471,985 | 132,619,644 | 42.4 | 10.0 |
| 1x HBSS | R2 | 863,126,588 | 610,087,746 | 500,187,621 | 464,002,407 | 82.5 | 17.2 |
| HEPES + 1mM EDTA, 1 h | R1 | 715,788,323 | 505,049,377 | 370,779,196 | 333,864,494 | 74.8 | 25.9 |
| Return, 1xHBSS after EDTA | R1 | 706,960,951 | 484,478,770 | 389,402,191 | 350,509,296 | 76.4 | 19.1 |
| 1x HBSS | R3 | 128,374,188 | 100,671,274 | 66,707,754 | 63,677,166 | 68.8 | 33.4 |
| HEPES + 1mM EDTA, 1 h | R2 | 116,329,893 | 85,695,816 | 47,499,071 | 43,718,226 | 42.1 | 42.7 |
| Return, 1xHBSS after EDTA | R2 | 162,888,591 | 123,015,944 | 95,068,131 | 90,774,223 | 68.7 | 21.9 |
